# Clonal dynamics deviate from neutral drift in zebrafish spermatogenesis

**DOI:** 10.64898/2026.06.08.730911

**Authors:** Jenna M. Weber, Connor R. Shrader, Drew B. Shapiro, Kimberly T. Truong, Andrea L. Sposato, Frederick R. Adler, James A. Gagnon

## Abstract

Spermatogonial stem cells (SSCs) maintain male fertility, but how their clonal dynamics change with age remains poorly understood. Here, we use in vivo CRISPR barcoding in zebrafish to label SSCs and track their contributions to sperm production through monthly sampling across the fertile lifespan. We find that only a subset of embryonic germ cells contributes to adult sperm production and that the relative contributions of individual clones shift dramatically over time. To interpret these dynamics, we developed a mathematical model that quantifies clonal drift rates and formally tests neutral versus non-neutral hypotheses. Most SSC clones show significant evidence of drift, yet their dynamics are inconsistent with a neutral model where clones have equal fitness and their dynamics are driven solely by random chance. Instead, we observe heterogeneous clonal dynamics, including a positive relationship between clone size and drift rate. Together, these findings demonstrate that SSC clonal dynamics are non-neutral across the zebrafish reproductive lifespan with important implications for allele transmission.

## INTRODUCTION

Adult stem cells sustain tissue integrity throughout life even as their populations undergo continual remodeling. Across different tissues, stem cell dynamics are governed by stochastic turnover and neutral competition between clonal lineages that each descended from a distinct ancestral stem cell. As a result, stem cell clones expand or contract over time by clonal drift (Klein & Simons, 2011).

Tissue architecture, proliferation rates, and extrinsic signals from the stem cell niche can shape the pace and pattern of stem cell clonal dynamics (Chatzeli & Simons, 2020; O’Brien, 2022; Rando et al., 2025; Ritsma et al., 2014). Moreover, mutations that accumulate with age can lead to biased competition between clones with potentially negative consequences to the organism, including clonal hematopoiesis and cancer (Bowling et al., 2019; Kapadia & Goodell, 2024; van Neerven & Vermeulen, 2022).

Germline stem cells are uniquely important because competition between clones can drive biased transmission of alleles to subsequent generations (Krieger & Simons, 2015; Lehmann, 2012; O’Brien, 2022; Weissman, 2015). In the male germline, spermatogonial stem cells (SSCs) within the testis sustain continuous sperm production. Maintaining lifelong fertility without depleting the SSC population requires a balance of differentiation and self-renewal. Various strategies have evolved in animals to maintain this balance. In *Drosophila melanogaster*, physical contact with the somatic hub stem cell ensures that SSCs divide asymmetrically (Fuller & Spradling, 2007). One daughter cell retains contact with the hub and maintains SSC identity while the other loses contact and differentiates. This simple constraint elegantly balances stem cell maintenance and a stable production of sperm throughout adulthood. By contrast, vertebrates rely on population asymmetries to maintain germline stem cells. A pioneering study of the mouse germline used single-clone labeling to suggest that most SSCs obey neutral drift dynamics, in which all SSCs have an equal chance to divide or differentiate and clone sizes grow or shrink stochastically (Klein et al., 2010; Klein & Simons, 2011). Emerging germline data from other systems, including human data, show that positive selection plays an important role in shaping stem cell dynamics in the testis (Goriely & Wilkie, 2012; Kanatsu-Shinohara et al., 2016; Neville et al., 2025; Seplyarskiy et al., 2025). While neutral drift has been a useful framework for understanding stem cell dynamics, increasing evidence from lineage tracing and human germline studies indicate that clonal behavior may be influenced by heterogeneity and selection.

Our understanding of SSC behavior in vivo has been limited by the challenge of tracking individual clones across the fertile lifespan. Physiological SSC population dynamics cannot be easily modeled outside of the organism (Kaltsas et al., 2025), and transplantation of labeled clones inherently perturbs the testicular niche (Lord & Oatley, 2018; Nóbrega et al., 2010; Ogawa et al., 1997). Both of these methods add confounding selective pressures to the SSC pool that may interfere with endogenous dynamics. Furthermore, traditional clone labeling tools generate only a small number of distinguishable labels, so homoplasy, where two clones are identically labeled, can obscure conclusions (Kanatsu-Shinohara et al., 2016; Kawahara et al., 2025; Klein et al., 2010). These methods also often rely on tamoxifen-induced recombination, which disrupts endocrine signaling and can cause adverse effects in the developing testis (Patel et al., 2017; van der Ven et al., 2007). Furthermore, visualizing labeled stem cells within the testis is an endpoint assay, making repeated observation of the same population impossible (Kawahara et al., 2025; Nguyen et al., 2020).

Sampling sperm provides a non-destructive method to observe the progeny of a stem cell population and thereby infer stem cell population dynamics. However, fluorescent labels generated in SSCs are often dim or absent in differentiated sperm, and serially sampling sperm from mammalian model organisms is labor intensive (Anciuti et al., 2025). Therefore, understanding SSC clonal dynamics in vivo requires a new research model that can simultaneously trace sperm lineages and quantify parameters of clonal expansion across adulthood.

Classical mutagenesis studies revealed that the zebrafish germline is mosaic, with distinct embryonic germ-cell lineages contributing unequally to the adult germline (Mullins et al., 1994; Solnica-Krezel et al., 1994; Walker & Streisinger, 1983). More recently, CRISPR-based approaches demonstrated that individual founders can transmit multiple independently edited alleles through the germline, confirming substantial germline mosaicism at nucleotide resolution (Brocal et al., 2016; Höijer et al., 2022; Hwang et al., 2013). Likewise, CRISPR lineage-recording studies showed that multiple embryonic germ-cell lineages contribute to the gamete pool (Junker et al., 2017). Together, these studies establish that the zebrafish germline is clonally heterogeneous and that multiple embryonic lineages contribute to adult gamete production. However, whether the relative contributions of those lineages remain stable or fluctuate over the reproductive lifespan is unknown. Zebrafish are uniquely suited to address this question because sperm can be repeatedly sampled from the same individual without sacrificing the animal, enabling direct observation of germline output over time.

Here, we combine multiclonal in vivo CRISPR barcoding with longitudinal sperm sampling to reconstruct SSC population dynamics across the fertile lifespan of zebrafish. We then interpret these longitudinal data by developing a novel mathematical model that incorporates the structure of our tracing system, the temporal nature of the dataset, and the uncertainty inherent in inferring SSC clone size from sperm output. We find evidence that clone sizes change over time but the dynamics do not conform to a “neutral” hypothesis that assumes all clones renew at the same rate and with equal fitness. Instead, we propose that heterogeneity in clonal dynamics arises through genetic or spatial variation in the testis. Overall, our combined experimental and mathematical strategies reveal how stem cell populations change in the native context of the aging zebrafish testis.

## RESULTS

### Modifying GESTALT barcoding to study clonal dynamics in the testis

Over the past decade, CRISPR tools have emerged for cell lineage tracing in animal models (Askary et al., 2025; Masuyama et al., 2022; McKenna & Gagnon, 2019; Sankaran et al., 2022). In these approaches, CRISPR editing of genomic barcodes generates a diverse set of heritable “barcodes” that can be recovered and used to classify cells into pedigrees. However, the DNA barcodes generated by these methods have mainly been recovered from dissected organs and dissociated cells from endpoint assays. We hypothesized that these methods could be adapted to measure germline clonal dynamics through repeated germ cell sampling from the same individual.

We developed a strategy to label primordial germ cells (PGCs) in developing zebrafish with CRISPR barcodes using GESTALT lineage tracing (McKenna et al., 2016; Raj, Wagner, et al., 2018) (Fig. 1A). Our approach harnesses CRISPR editing of the GESTALT barcode at two developmental stages following existing protocols (Raj, Gagnon, et al., 2018). In zebrafish, four cells inherit germ plasm and are specified as PGCs during the first four hours of development (Raz, 2003; Weidinger et al., 1999; Yoon et al., 1997). We applied the first round of barcode editing to cover this time period by microinjecting 1-cell embryos with Cas9 protein and barcode-specific gRNAs. During and after gastrulation, the PGCs migrate to the future location of the gonad, dividing to generate ∼30 cells by 24 hours post-fertilization (hpf) (Tzung et al., 2015). Thus, we applied a second round of barcode editing at 24 hpf using a heat shock-inducible Cas9 transgenic line, which engaged a second set of gRNAs to complete GESTALT barcode editing. The number of PGCs at 24 hpf remains relatively stable during the first week of development when somatic gonad formation begins (Leerberg et al., 2017; Tzung et al., 2015). Domesticated zebrafish lack sex chromosomes and maintain a bipotential gonad until approximately 25 days post-fertilization (dpf) when the immature gonad starts to adopt either an ovary or testis fate (Chen & Ge, 2013). The number of PGCs that survive this transition to sex-specific germline stem cells remains unclear (Barton et al., 2016; Bertho et al., 2021). Zebrafish reach sexual maturity between 3-4 months of age (Liew & Orbán, 2014). Our animals were grown to adulthood and males were selected for future experiments.

**Figure 1.**
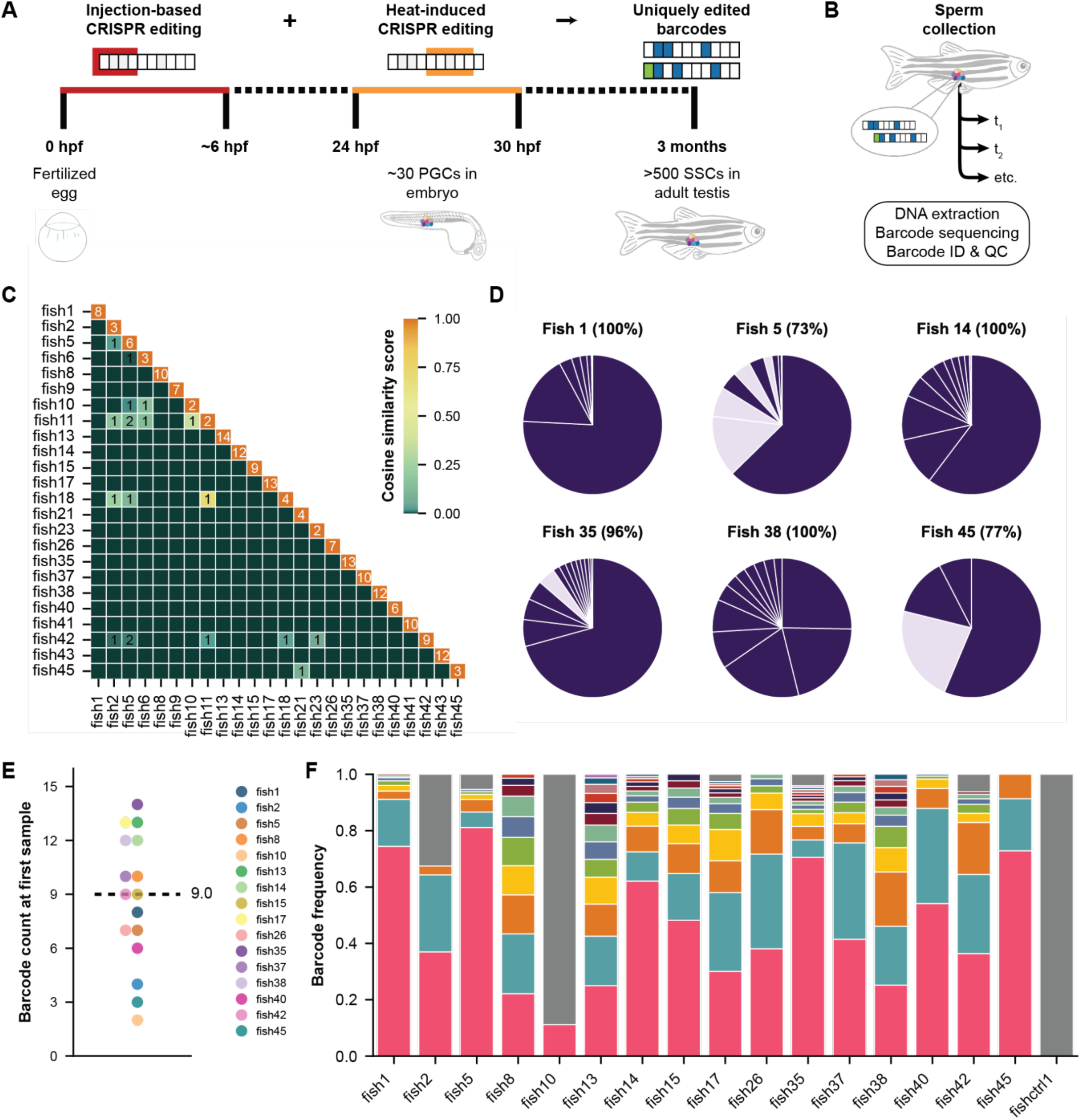
CRISPR barcodes that label germ cells can be recovered from sperm samples. A) For GESTALT barcoding of zebrafish germ cells, two rounds of CRISPR-Cas9 editing generate random insertions and deletions in the barcode at key time points in zebrafish gonad development. B) Barcode gDNA from sperm samples is amplified for sequencing, barcode identification, and processing. C) Heat map comparing barcode distributions between fish using cosine similarity score. Diagonal entries count unique barcodes within fish; off-diagonal entries count shared barcodes between fish, if >0. D) Pie charts depicting barcodes unique to each fish (dark slices) and barcodes found in multiple fish (lighter slices). Percentage of unique reads that were kept for later data analysis (sum of all darker slices) noted above each pie chart. E) Count of unique edited barcodes recovered from each fish after the first sperm sample. Each dot represents the count for that fish across all three technical replicates. Black line and numeral notate the median across all fish (9, range 2 to 14). F) Quantification of all barcode frequencies in each fish’s first sperm sample. The same color scheme is used for each fish even though their barcodes are unique. Gray represents the unedited barcode.

### GESTALT barcodes can be recovered from sperm

In male animals, embryonic PGCs give rise to spermatogonial stem cell clones within the testis which will subsequently generate sperm. In male zebrafish, PGCs give rise to approximately 500-1000 SSCs in adults (Mullins et al., 1994). The mapping of PGCs to SSC clones remains a mystery. To confirm that the barcodes generated in the PGCs are faithfully inherited through differentiation and meiosis, we sampled sperm from adult males (Draper & Moens, 2009) beginning at 5 months of age and measured barcodes from sperm genomic DNA using amplicon sequencing (Fig. 1B).

Sequencing reads were translated into barcodes using a modified GESTALT pipeline with a custom quality-control filtering process optimized for sperm samples (see Methods, Fig. S1). Sperm samples were processed in technical triplicate at each time point, and strong concordance between replicates was observed for all samples (Fig. S2A). These results demonstrate that GESTALT barcodes are stably transmitted through the germline and reliably detected in the sperm, validating their use as heritable lineage markers for SSC differentiation.

We generated a cohort of barcoded adult males for longitudinal sperm sampling, along with four unedited males to serve as controls (i.e. males carrying the barcode but did not undergo CRISPR-Cas9 editing). After the first sperm sampling time point at five months of age, we compared the barcodes between individuals to assess how well the method discriminates between animals.

Barcodes are remarkably unique to each animal. Cosine similarity scoring demonstrates that most fish have completely distinct barcodes (Fig. 1C). On average, only 2.7% of barcodes were recovered from two or more fish (Fig. 1D; Fig. S3). These barcodes were removed for subsequent analysis unless clear evidence supported retaining them in one fish (see Methods).

We then counted the barcodes present in each sample. In the edited animals, a median of 9 edited barcodes were recovered per fish (range 2 to 14) (Fig. 1E), lower than the 30 predicted by known PGC numbers during early zebrafish development (Raz, 2003). This discrepancy may reflect loss of PGCs before the SSC stage, failure of some SSCs to generate sperm, or incomplete CRISPR-Cas9 editing across the PGC pool. Nevertheless, the number of recovered barcodes was sufficient to resolve the clonal contributions of individual SSC lineages within each animal, establishing the baseline editing efficiency and barcode diversity in our cohort.

Interestingly, each sperm sample contained a mixture of barcodes at various abundances, showing that some labeled SSC clones were contributing more to sperm production than others at the time of sampling (Fig. 1F). Although the unedited barcode appeared in 6 out of 17 edited fish, on average it only represented 23% of the barcode population in those fish (range 3-89%). As expected no edited barcodes appeared in the sperm collected from the unedited control animal. Together, these data and previous studies (McKenna et al., 2016; Raj, Wagner, et al., 2018) demonstrate that in vivo CRISPR barcoding with GESTALT can generate a diverse set of immutable labels that uniquely label progenitor cells and their descendants with higher confidence than alternative methods (Ikeda et al., 2025; Pei et al., 2017).

### Longitudinal samples demonstrate how clonal contributions to sperm production fluctuate with time

Having confirmed our ability to recover barcodes from sperm, we continued sampling the cohort monthly across their fertile lifespan (Fig. 2A). Peak fertility in zebrafish occurs between 7 and 18 months of age (Westerfield, 2000) and fertility declines in older zebrafish (Sposato et al., 2024). Laboratory zebrafish are generally retired after 24 months of age. Our sampling scheme covers these major time points, beginning at 5 months of age and ending when the fish died or reached 26 months of age. Sampling was not always successful — some males occasionally failed to produce sperm at the time of sampling. Of the 49 male fish in our cohort, only six provided sperm at every time point but 26 fish provided samples at least half of the time. Some fish died of natural causes and dropped out of our cohort, at similar rates to control animals which were edited but not sampled. While sampling of younger males was occasionally detrimental to their survival, sampling throughout adulthood had little impact on survival. This longitudinal sampling paradigm is uniquely possible in teleost fish like zebrafish, which are large enough for non-lethal sperm recovery (Draper & Moens, 2009) but do not require complex sampling methods like in mammals.

**Figure 2.**
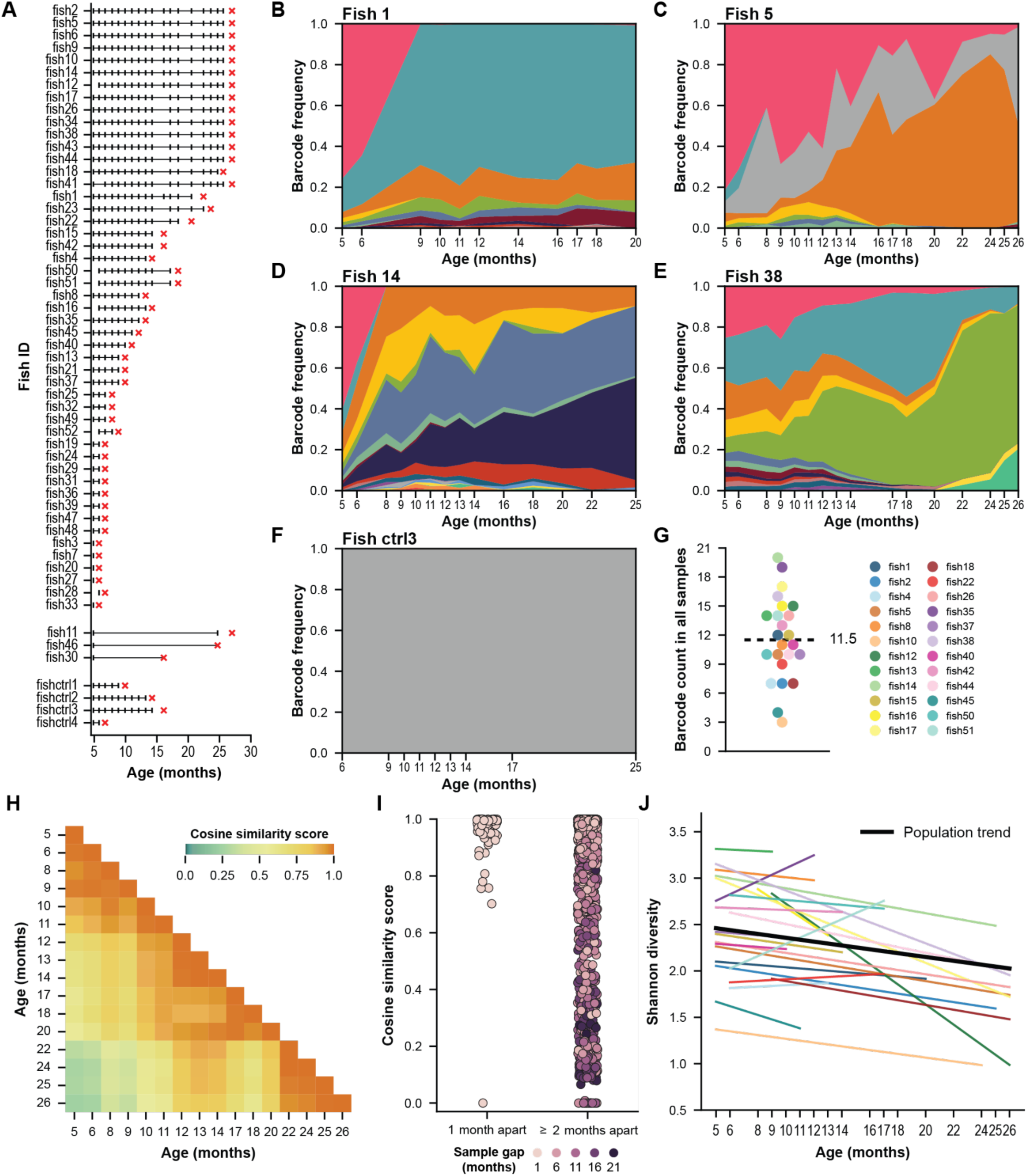
A consistent set of clones generate sperm across the zebrafish lifespan. A) Longitudinal sperm sampling timeline. Each fish is represented by a horizontal line showing the period of continuous observation, and tick marks show sampling time points that produced at least some visible volume of sperm for that individual. Red x’s indicate age at death. Lines are ordered by the number of successful samples. Data began at age 6 months for males who could not be reliably sexed at 5 months of age. Small gaps in the timelines of certain individuals represent unsuccessful sampling (no volume of sperm produced by abdominal massage). B-F) Barcode frequencies across time for five example individual fish. The same color scheme is used for each fish even though their barcodes are unique. Gray represents the unedited barcode. G) Count of total unique edited barcodes recovered from each fish over the sampling period. Each dot represents the count for that fish across all three technical replicates of all sperm samples. Black line and numeral indicate the median across all fish (11.5, range 3 to 20). H) Heat map comparing barcode distributions between all sperm samples of Fish 38. Color determined by cosine similarity score for each pairwise comparison. I) Summary of cosine similarity scores between samples for all fish. Each point represents a pair of samples from a single fish. Color determined by the number of months separating the two samples. J) Shannon diversity index across time for each fish. Each colored line was fit using a linear mixed model, see G for legend. Black line represents the population trend.

Using our longitudinal sperm barcode data, we visualized changes in clonal proportions across the fertile lifespan for each fish (Fig. 2B-F; Fig. S4). Again, technical replicates demonstrated striking similarity throughout the sampling period (Fig. S2B-C). We were surprised to discover substantial variability in how SSC clones contributed to sperm production over time. Some clones initially dominated the pool but then disappeared from all later samples (Fig. 2B, pink clone), while others contributed very little to sperm production at early reproductive ages and later expanded their output dramatically (Fig. 2C, orange clone). Other clones remained remarkably consistent across many months (Fig. 2D, red clone). New clones rarely emerged at late reproductive ages (Fig. 2E, pale yellow and dark green clones) and no edited clones ever appeared in the unedited control animals (Fig. 2F). Overall, these visualizations suggest that clonal activity varies both across time and between individual fish.

However, the total number of clones that ever contribute to sperm production (median = 11.5 clones) is only slightly larger than the number of clones detected in the first sample (median = 9 clones) (Figs. 1E, 2G). Taken across all fish, samples from consecutive months were more likely to be similar than samples separated by longer time frames, showing that clonal proportions are shifting incrementally over time (p < 10^-27^, Kolmogorov-Smirnov test) (Fig. 2H-I; Fig. S5). This demonstrates that a consistent set of SSC clones contributes to sperm production across the zebrafish lifespan but that their contributions fluctuate with time.

### Computational modeling to understand clonal dynamics

The serial sperm samples revealed that the relative contribution of each SSC clone to the sperm pool varies over time. This variation could reflect biological features of clonal dynamics, such as neutral drift, selection, or clone-dependent heterogeneity in sperm output. However, it may also be influenced by sampling error and technical noise from library preparation for sequencing. Under neutral drift, clonal diversity within a population decreases over time (Gillespie, 2004). Using the Shannon diversity metric, we found that clonal diversity decreased with age in most individual fish. However, a linear mixed model indicates that the overall diversity across all fish does not show significant evidence of decreasing over time (Fig. 2J; Fig. S6). Interestingly, fish with increasing diversity are typically characterized by a single disproportionally large clone at the initial sample that later shrinks (fish 35 & 51, Fig. S4, S6). To better quantify characteristics of these observed dynamics, we developed a computational model of stem cell clonal dynamics, sperm differentiation, and serial sperm sampling.

Our mathematical model uses a modified Moran process to describe SSC population dynamics and sperm production (Moran, 1958). The Moran process is a stochastic model of population genetics that has previously been adapted to model stem cell dynamics in other contexts (Clayton et al., 2007; Lopez-Garcia et al., 2010; Sun & Komarova, 2012). In our model, each individual stem cell undergoes random and spontaneous division at rate *r* to produce sperm (Fig. 3A). Each division has probability *p* of being a symmetric stem cell differentiation, which removes the stem cell and produces sperm. To maintain a constant stem cell population, this differentiation is coupled with the symmetric division of another stem cell, potentially from a different clone. This allows individual clones to increase and decrease in size over time even if all cells are equally capable of proliferating. All stem removal is through differentiation; we do not explicitly model stem cell death.

**Figure 3.**
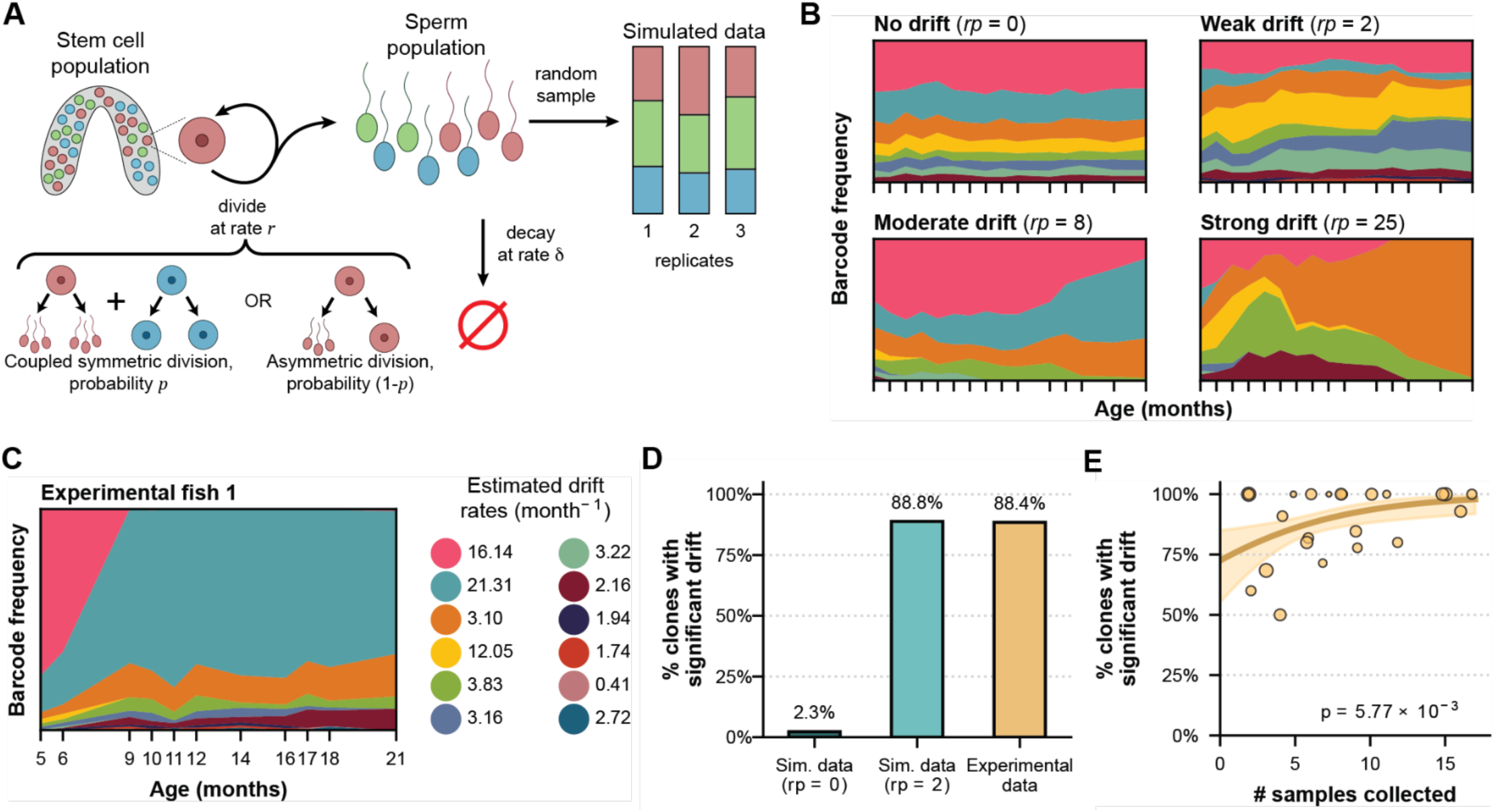
Experimental clones show strong evidence of drifting. A) Diagram of a mathematical model that represents neutral clonal dynamics and sperm sampling. The model allows each individual stem cell to randomly divide or differentiate at rate *r*. Cells divide asymmetrically with probability 1-*p* or differentiate with probability *p*. To maintain a constant SSC population size, stem cell differentiation induces another stem cell to divide symmetrically. The rate of symmetric division, *rp*, represents the overall rate of drift. B) Sample model simulations with different rates of drift. C) Differentiation rates for each clone are inferred from clonal data using maximum likelihood estimation on the model. D) Percentage of simulated and experimental clones where significant evidence of drift is detected using the maximum likelihood method. E) Relationship between the lifespan of experimental fish and the percentage of experimental clones with significant evidence of drift. Each point represents a fish and the point size corresponds to the number of clones in the fish. A logistic model fit shows significant correlation between the number of sperm samples collected from the fish and the proportion of clones identified as having non-zero drift (p=5.77 x 10^-3^).

We also do not consider quiescent stem cells, since such cells would not produce sperm. The rate that differentiations occur (*rp*) represents the strength of neutral drift in the stem cell population.

Model simulations show that increasing this parameter creates clones that change more rapidly (Fig. 3B). Stem cells may also divide asymmetrically with probability (1-*p*), which produces sperm without altering the stem cell population. Sperm naturally decays at rate δ until sampling. Noise was added to simulated samples to mimic the technical noise introduced by experimental sampling and replicate library preparation.

We determined model parameter values from past studies on zebrafish spermatogenesis. The cell cycle rate *r* was set to ∼36 per month (one cycle per 20 hours) (Nóbrega et al., 2010) while the sperm removal rate δ was estimated as ∼6 per month, equivalent to a mean sperm lifespan of 5 days (Cattelan & Gasparini, 2021) (see Methods).

### Most clones show significant evidence of drift

We first used our model to evaluate whether the experimental data are consistent with a model with unchanging clone sizes (*rp* = 0). Maximum likelihood estimation was used to compute the rate of neutral drift, *rp*, most likely to reproduce the observed data in each experimental clone. In Fish 1, for instance, clones have measured drift strengths ranging from 0 to 21 differentiations per month (Fig. 3C). This estimator showed little bias on simulated data, making it suitable for use on experimental data (Supplemental Text). A likelihood ratio test was used to determine whether the maximum likelihood estimate for *rp* is significantly more likely than the likelihood of unchanging clone sizes (Zar, 2014) (see Methods). Our method is conservative; in simulated fish with no drift, only 2.3% of clones were falsely identified as undergoing drift (Fig. 3D). In the experimental data, 88.4% of experimental clones have significant evidence of drift. This indicates that the temporal changes in clone size in our data cannot be explained by a model of unchanging stem cell populations.

We next tested to what extent our statistical test depends on the amount of available clonal data using a logistic regression model. A significant positive correlation was observed between the number of collected samples in experimental fish and the evidence of drift in its clonal data (Fig. 3E, p = 5.77 x 10^-3^). As expected, drift detection depends on multiple samples over time from a given individual, suggesting that our estimate of the fraction of clones undergoing drift may be conservative. These results indicate that the vast majority, perhaps all, of SSC clones are not fixed in size in adult zebrafish but instead exhibit temporal dynamics.

### Neutral drift is insufficient to explain experimentally-observed clonal dynamics

Next, we evaluated whether the experimentally-observed clonal data conform to a “neutral hypothesis” in which all clones are equally capable of proliferating. Our model assumes neutrality by giving clones equal cell cycle rates. Comparison between the quantitative predictions of the model and the observed clone tracing data revealed several discrepancies that demonstrate that SSC clones in the zebrafish testis are not drifting neutrally. Under a neutral model, stem cell differentiation rate should not depend on clonal identity. Using our likelihood method, we computed the rate of drift in all the observed clones and compared this distribution to the drift strengths measured from simulated data. To enable direct comparison, the initial clone distributions and sample times of simulated fish were set to match the experimentally-measured clones. The median drift rate estimate in the measured clones is 2.16, so we set *rp=2* in simulations. If the observed clones share a fixed drift rate, then the measured drift rates will have the same variance as drift rates measured from the simulated fish. Instead, the drift rate of observed clones have a much wider distribution than simulated clones (Fig. 4A). This suggests that SSC clones have different effective rates of drift, rather than a single inherent rate.

**Figure 4.**
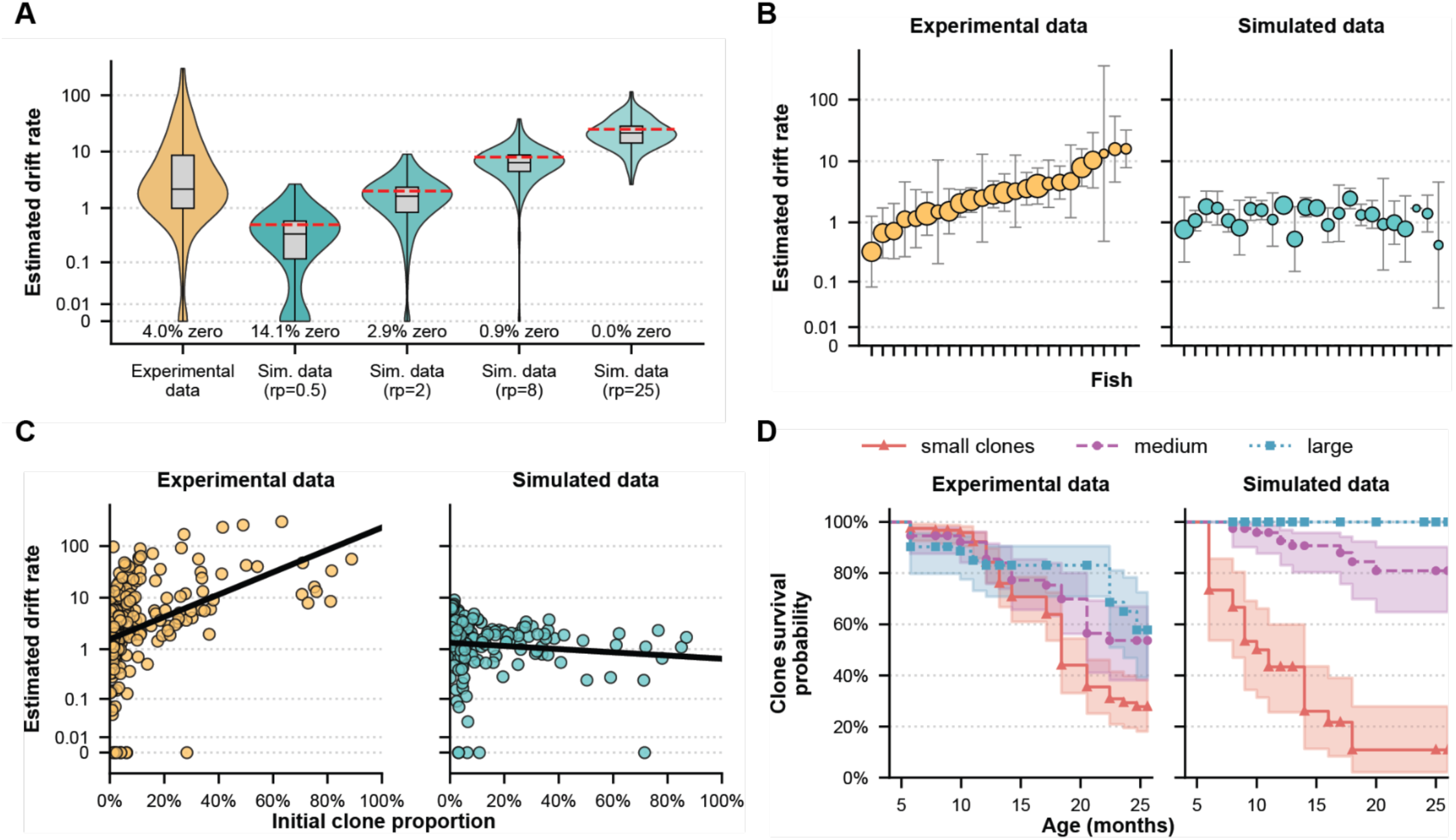
Observed clonal dynamics do not conform to neutral drift. A) Distribution of symmetric division rate (*rp*) estimates in simulated and experimental data. True drift rates for the simulated fish are shown in red dashed lines. B) Mean and variance of drift estimates in each simulated and experimental fish. Fish are ordered so that means are increasing in the experimental fish. C) Correlation between initial clone size and estimated differentiation rate. Each point represents one clone from one fish and the black lines are the best linear fits. D) Clone survivorship in simulated and experimental data. Small clones are defined as having initial proportion <5%, medium clones 5-30%, large clones >30%.

To assess whether clonal drift rates vary between fish while remaining consistent within individuals, an overall drift rate was computed for each simulated and experimental fish using the geometric average drift rate across its clones. Fish-averaged drift rates have greater variance in the experimental fish when compared to simulated fish (p=2.15 x 10^-3^, Levene test, Fig. 4B), indicating that some fish indeed have higher overall drift rates than others. Within-fish variance in drift strength was then quantified by measuring the residual drift strength of each clone relative to the fish-level mean. This within-fish variance in experimental fish is significantly greater than in simulated fish (p=1.42 x 10^-6^, Levene test). Together, these results indicate that the observed heterogeneity in clonal drift rates is only partially explained by differences between individual fish.

Under the neutral hypothesis, stem cell behavior is independent of clone size. However, we observed that many of our fish contain small SSC clones that drift slowly and large clones that change more rapidly (examples in Fig. 2B-E, S4). To test whether this pattern reflects a systematic relationship, a linear regression model was fit with drift strength as a function of initial clone size, with random effects for each fish. A significant correlation between drift strength and initial clone size was observed in the experimental data but not in simulated data (p=2.70 x 10^-12^ and p=0.176, respectively) (Fig. 4C, see Methods). This relationship is not predicted by the neutral model and shows that sperm production is heterogeneous between clones.

The dependence of clonal behavior on clone size is further supported by survivorship analysis. Under a neutral hypothesis, small clones are more vulnerable to elimination than large clones. Stratification by initial clone size showed that this dependence of clone survival on initial size is present in the experimental clones, but is markedly weaker than predicted by the model (p=8.31 x 10^-9^, Weibull AFT interaction, see Methods, Fig. 4D). Clones of different initial sizes therefore exhibit more similar risks of elimination than expected under a neutral model. Notably, small clones remain relatively stable when compared to large clones, suggesting that the factors driving heterogeneous clonal dynamics are linked to overall sperm production. These findings were robust to a range of barcode processing and simulation parameters (Fig. S7 and see Methods). Together, these findings indicate that SSC clones exhibit diverse behaviors that are incompatible with neutral drift, and that this clonal heterogeneity is only partially explained by differences between fish.

## DISCUSSION

In this study, we combined in vivo GESTALT barcoding with longitudinal sperm sampling to quantify SSC clonal dynamics across the zebrafish reproductive lifespan. These data reveal that SSC clones do not remain stable in size and do not follow neutral drift dynamics. Instead, clonal contributions to sperm production are dynamic and heterogeneous over time. By integrating lineage tracing with a quantitative model of stem cell dynamics, we are able to formally test and reject neutral hypotheses rather than relying on descriptive inference alone. Clones display a wide range of drift rates, with larger clones tending to change more rapidly than smaller clones. In addition, initial clone size is only a weak predictor of long-term persistence, as many large clones are lost while smaller clones remain stable. Together, these observations indicate that SSC clones make unequal effective contributions to sperm production, revealing structured heterogeneity in clonal behavior that cannot be explained by neutral competition alone.

Several mechanisms could cause such non-neutral dynamics. Intrinsic differences between clones such as lineage-specific gene expression programs or acquired mutations could bias proliferation or differentiation rates. In other stem cell systems, mutations that accumulate with age can drive biased clonal expansion and alter competition dynamics (Bowling et al., 2019; Kapadia & Goodell, 2024; Klein & Simons, 2011; van Neerven & Vermeulen, 2022). These theories do not fully explain our observations; notably, that the most expanded clones found in the initial sperm sample from young fish are also highly susceptible to elimination in later samples. Alternatively, extrinsic factors including unequal access to niche signals and systemic physiological changes may differentially regulate SSC activity (Cui et al., 2025; O’Brien, 2022; Rando et al., 2025). Spatial organization within the testis may also play an important role. The distribution of clones along seminiferous tubules in the testis likely creates nonuniform local constraints on clonal competition. At the organ scale, the two testis lobes may be asymmetric in their sperm production. Our data do not distinguish between these possibilities but instead indicate that one or more of these factors generate persistent heterogeneity in SSC behavior across the lifespan.

Our findings contrast with prior studies of SSC dynamics in mice. Pulse-labeling experiments in mouse testes have supported neutral drift, with clone survival decreasing over time and surviving clones expanding in size (Klein et al., 2010). More recent work has suggested that clonal contributions are stable on the testis-wide scale despite the local turnover due to population size effects and geometric considerations (Ikeda et al., 2025). Several factors may explain the differences between these studies and our results. First, mouse studies of germ cell dynamics often focus on spatially localized regions of seminiferous tubules, whereas our approach captures testis-wide dynamics. Second, our longitudinal sampling spans the full reproductive lifespan, potentially revealing temporal variability not captured in shorter experiments. Finally, species-specific differences in testis organization or SSC regulation may contribute to fundamentally distinct clonal dynamics between zebrafish and mammals (Schlatt & Ehmcke, 2014; Yoshida, 2016).

Our study has several limitations. Most importantly, sperm output provides an indirect measurement of SSC population dynamics, and changes in clonal representation may reflect differences in differentiation output rather than underlying stem cell number, although previous studies in zebrafish suggest that stem cell number remains relatively stable with age (Sposato et al., 2024). In addition, our approach does not capture spatial information, limiting our ability to distinguish between intrinsic and niche-driven mechanisms. Technical constraints including low sperm volume (Hagedorn & Carter, 2011) and occasional sample loss reduce the resolution of some time points and may limit sensitivity to subtle changes in clonal behavior. These factors likely make our estimates of drift conservative and highlight the need for complementary approaches that directly measure SSC populations and their spatial organization.

Despite these limitations, our work establishes a framework for studying clonal dynamics in the vertebrate germline across the fertile lifespan. The observation that SSC populations exhibit non-neutral dynamics has important implications for how genetic variation is transmitted across generations, as unequal clonal contributions could bias allele transmission over time (Krieger & Simons, 2015; Weissman, 2015). Consistent with that idea, age-associated clonal expansions of spermatogonial stem cells carrying specific mutations in the human germline have been linked to biased transmission and increased risk of disease, a phenomenon often referred to as selfish spermatogonial selection (Choi et al., 2008; Goriely & Wilkie, 2012; Maher et al., 2018). Our results establish a tractable in vivo system for quantifying such dynamics and provide a foundation for testing how aging, niche perturbations, and disease-associated mutations influence stem cell competition. Future studies integrating lineage tracing with spatial assays and functional manipulations will be critical for identifying the mechanisms that drive heterogeneous behavior in SSC populations.

## METHODS

### Zebrafish husbandry

All zebrafish used in this study were housed at the University of Utah CBRZ facility. This research was conducted under the approval of the Office of Institutional Animal Care and Use Committee (IACUC protocol 21-02017) of the animal care and use program of the University of Utah. Male fish used for sperm sampling were housed in 2L tanks and separated with clear dividers to maintain social interaction while allowing for correct re-identification of individuals and preventing the loss of sperm to background breeding.

### CRISPR barcoding

A new single-copy transgenic line of the GESTALT barcode containing an array of 10 CRISPR target sites, with minor modifications to the flanking sequences was generated as previously described (Raj, Gagnon, et al., 2018) and given the line designation zj10Tg. Double transgenic animals were generated by crossing the GESTALT barcode line to an inducible Cas9 line as previously described (Raj, Gagnon, et al., 2018). Briefly, transgenic females containing one copy of the GESTALT barcode sequence^zj10Tg^ were crossed to Tg(*hsp70l*:zCas9-2A-EGFP, 5×(U6:sgRNA))^a168Tg^ males. The resulting embryos were injected at the one-cell stage with 1 nl of the following injection mix: 2 µl 1 M KCl (Sigma-Aldrich), 1 µl phenol red (Sigma-Aldrich), 1 µl SpCas9 protein (IDT), and 2 µl sgRNA mix at molar ratios (200 ng/uL each). Injected sgRNAs are specific to sites 1-4 of the GESTALT barcode array. These embryos were screened for green hearts to identify the barcode transgene and then heat shocked at 24 hpf to induce SpyCas9 expression and engage a second set of sgRNAs specific to sites 5-9 of the GESTALT barcode array. Double transgenic embryos were identified by GFP expression in the whole body at 48 hpf and were grown to adulthood.

### Sperm sampling and genomic DNA extraction

Starting at 5 months of age, sperm collection was performed as previously described (Draper & Moens, 2009). Briefly, males were anesthetized with 0.02% Tricaine (MS-222) and placed belly up on a moist sponge under a Leica S9i stereomicroscope, illuminated with a 6 Watt LED Dual Gooseneck Illuminator (AmScope). The anogenital area was gently dried with a KimWipe to avoid contamination. Sperm were elicited by gently squeezing the abdomen with smooth-sided forceps and collected using a 5-uL glass microcapillary (Drummond Scientific Company cat #2-000-001). Immediately after collection, sperm were released into 10 uL of ultrapure water and maintained on ice until genomic DNA extraction using a modified HotSHOT method (Meeker et al., 2007) (2 uL 6x lysis buffer, 3uL 2.5x neutralization buffer). Following the sperm sampling procedure, fish were returned to their home tank. Fish were sampled once every four to six weeks.

### Amplicon sequencing of sperm barcodes

Barcode sequencing libraries were prepped in triplicate from sperm genomic DNA. Three aliquots of a sperm sample, each containing up to 500 ng of gDNA, were loaded into separate 25 μl PCR reactions and amplified using primers immediately flanking the barcode (Gagnon et al., 2018; McKenna et al., 2016; Takasugi et al., 2022). If there was less than 1500 ng from a sample, it was split evenly between the three reactions. The second PCR added dual sequencing indices unique to each sample following published protocols (Takasugi et al., 2022). PCR products were purified with Mag-Bind® TotalPure NGS magnetic beads (Omega Bio-Tek cat #M1378-01) after each reaction. Amplicons were pooled at roughly equimolar ratios and sequenced using Illumina MiSeq Reagent Kits v2 (500 cycle).

### Computational analysis and data quality control

Sequencing data were processed using the previously published GESTALT pipeline, with modifications to permit the use of Singularity as a container environment on our computing cluster (McKenna et al., 2016; Takasugi et al., 2022). We set the GESTALT trim quality and match proportion parameters to 10 and 0.8, respectively. Reads are trimmed where the average quality score in a sliding window falls below the defined trim quality parameter. Listing base pairs from left to right, only base pairs to the left of the sliding window are kept. Paired-end reads are discarded when the match rate of the overlapping section falls below the specified match proportion parameter.

Raw GESTALT outputs were refined using quality control parameters, including: 1) ≥30% of the reads in each sample FASTQ must be usable for the GESTALT pipeline and there must be ≥1000 of those reads. 2) Each barcode in a sample must be assigned ≥10 reads and represent ≥0.5% of the total reads in that sample. 3) For each sample, barcode sequences with Levenshtein distance ≤1 were connected by an edge to form a network. Connected components of this network were then collapsed to the most common barcode. 4) Each barcode in a sample must appear in ≥2 time points and in all three replicates at ≥1 time point. 5) If a barcode was detected in more than one fish, it was assigned to the fish where it had the highest total read count across all time points. To be assigned to a specific fish, the barcode needed to have at least 50X more reads in that fish than in all other fish combined. Otherwise, it was removed from all fish. 6) Sufficient data must remain after all previous steps, described below. Any data not passing these criteria were thrown out for all subsequent visualizations and analysis. All code for postprocessing and analyzing barcodes is available at https://github.com/Gagnon-lab/weber-shrader.

In some fish, the clonal proportions at one time point differed significantly from neighboring time points. We considered sperm samples from these time points to be anomalous: more likely explained by sampling error than underlying biology. We aimed to find a biological basis for the removal of these samples for the last step of quality control.

There were two quantitative metrics that tended to correspond to these visual anomalies. 1) The GESTALT pipeline outputs the total number of input reads as well as the number of successfully used reads. A low proportion of used reads to input reads indicates that many were not processed by GESTALT due to poor read quality. Samples where this proportion fell short of 0.3 were discarded. 2) For several samples, many of the original barcodes defined by GESTALT processing were later filtered out by the quality control pipeline. When the proportion of remaining reads after quality control to initial post-GESTALT reads for a given sample fell below 0.3, that sample was discarded. The mean and standard deviation of this proportion were taken for the remaining samples in each fish, and any samples falling at least 1.5 standard deviations below the mean were discarded (Fig. S8).

We noted that changing the GESTALT parameters affects the number of reads filtered. In particular, the trim quality and match proportion parameters affect the number of barcodes with only one read, which we call “singletons” (Fig. S9). By increasing trim quality to 14 or match proportion to 0.95, we found that the results are nearly identical (Fig. S10).

### Cosine similarity and diversity analysis

Cosine similarity scores were calculated using barcode identities and their relative abundances within each fish, summed across technical replicates. For each fish, we evaluated whether the cosine similarity of any two samples is negatively correlated with the time between those samples. To test this, we used a one-sided Mantel test (Legendre & Legendre, 2012; Mantel, 1967). We defined *X* as the matrix of cosine distances between pairwise time point samples for a fish (cosine distance = 1 – cosine similarity) and *Y* as a matrix that has the same size of *X* and entries equal to the time between samples (for example, sample months 5 and 7 are two months apart, so the corresponding entry in *Y* is 2). The Mantel test evaluates correlation between *X* and *Y* by repeatedly reordering one matrix at random and measuring how often the correlation is greater in the reordered matrix than the original matrix. Some, but not all, fish showed a significant correlation (Fig. S5).

Clonal diversity was measured using Shannon diversity (Shannon, 1948; Zar, 2014). The dependency of diversity on time was fit to a linear mixed model with random effects for the intercept. The coefficient for time was –0.021 and is not significant (p = 0.09).

### Model design and parameterization

We used a modified Moran process to describe spermatogonial stem cell differentiation and replacement in the testis (Moran, 1958). This stochastic model allows neutral clones to change in size over time. Stem cells divide at a fixed rate *r* and may either differentiate symmetrically (with probability *p*) or asymmetrically (with probability 1–*p*). Stem cells that differentiate produce a fixed number *s_d_* of barcoded sperm after a time delay of *τ* months. Sperm decay at rate δ until sampling. To mimic technical noise, simulated sperm samples are taken as a small multinomial sampling of the total accumulated sperm. More details of the model design are included in Supplemental Methods.

Important biological parameters for this model include the number of stem cells *N*, cell cycle rate *r*, differentiation time *τ*, sperm created per division *s*_!_, and sperm decay rate δ. These values are all estimated from available data. Stem cell count is set to 1000 (Mullins et al., 1994; Nóbrega et al., 2010), Cell cycle rate is set to 36 cell cycles per month (Nóbrega et al., 2010). The time delay *τ* from SSC to spermatid is 6 days, or 0.2 months (Leal et al., 2009). The total sperm produced from a differentiating stem cell is 1262 (Leal et al., 2009). However, because sperm are haploid, only half of them inherit our CRISPR barcode. Therefore, we set differentiated barcoded sperm *s*_!_from one stem cell as half of 1262, or 631. Sperm decay rate is set to 6.28 month^-1^, which was found by using a simplified model of sperm decay and fitting the resulting curve to data from (Cattelan & Gasparini, 2021) (Fig. S11A). Each simulated fish has initial clone frequencies set to match the first sample from a corresponding experimental fish. More information is included in Supplemental Methods.

We define the “drift rate” of a clone to be the value of the model’s differentiation rate (*rp* month^-1^) that is most likely to reproduce the observed data. Define *L*(*rp*) as the likelihood of observing a given clone’s sampling data given a differentiation rate *rp*. Then the maximum likelihood estimate of *rp* is

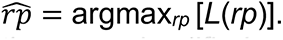

We perform maximum likelihood estimation on a simplified version of the model. The Supplemental Methods describes this method and shows that the method has low bias.

To compare model fits, we use the likelihood ratio test. Specifically, to determine whether a given clone’s data is consistent with the hypothesis that clone size does not change over time, we compute the test statistic

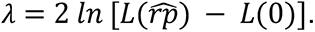

The model with a non-zero differentiation rate has one additional degree of freedom, so the level of 95% significance is 3.84 (Zar, 2014). Thus, we identify clones for which λ>3.84 as having statistically significant evidence of drift.

### Statistical tests

For the statistical tests further described below, most comparisons are made with simulations with *rp*=2. This is because the median drift strength estimated from our experimental data is 2.16.

We measured the dependence of drift evidence on the number of samples (Fig. 3E) using a logistic regression model. The coefficient of the initial clone frequency is 0.170 with significance p=5.77 x 10^-3^. We also fit the logistic regression to simulated fish with the same initial clone count and size and *rp*=2, repeated over ten independent simulation runs (not shown in the figure). The coefficient was significant with p<0.01 in all runs.

Between-fish drift strength (Fig. 4B) is measured by comparing the distributions of the mean symmetric-log drift strength (taken over all clones in a fish). We compared experimental fish to simulated fish with *rp*=2. Using the Levene test, we find that the experimental fish have more between-fish variance with p=2.15 x 10^-3^. Over nine repeated simulations, p<0.05 in eight of them, so this result is somewhat sensitive to the simulation outcome.

Within-fish drift strength (Fig. 4B) is measured by comparing the distributions of the residual symmetric-log drift strength (clone drift strength minus fish mean). We compared experimental fish to simulated fish with *rp*=2. Using the Levene test for equality of variance, we find that the experimental fish have more within-fish variance with p=1.42 x 10^-6^. We found p<10^-5^ in each of our nine repeated simulations.

Dependence of drift strength on initial clone size (Fig. 4C) is evaluated using a centered linear regression model predicting symmetric-log growth from initial clone frequency. We included random effects in the intercepts for each fish. The symmetric logarithm (with linear threshold at 0.01) is used because drift strength spans multiple orders of magnitude but can also equal zero. We find that drift strength does not significantly depend on initial size in the model simulations (p=0.176) but the interaction term for experimental data is significant (p=2.70 x 10^-12^). Over nine repeats of the simulations, p<10^-7^ in all of them.

Survivorship analysis is done using a Weibull-distributed accelerated failure time (AFT) model with predictors for initial clone size and clone type (experimental or simulated with *rp*=2). We use AFT instead of Cox proportional hazard because the Grambsch–Therneau test fails (Therneau & Grambsch, 2000). The coefficient of the initial frequency for the experimental data is 0.80 (p=0.0274). The coefficient of the interaction of model simulations with initial frequency is significantly higher (36.10, p=8.31 x 10^-9^). This interaction coefficient remained significant in nine repeated simulations.

These results are robust to the simulated drift rate being compared. In addition, these results do not significantly differ if we consider alternative data processing steps (see Fig. S7).

## AUTHOR CONTRIBUTIONS

Conceptualization: JMW, CRS, DBS, KTT, FRA, JAG

Data curation: JMW, CRS, DBS, ALS

Formal analysis: JMW, CRS, DBS

Funding acquisition: JMW, CRS, KTT, JAG

Investigation: JMW, KTT

Methodology: JMW, CRS, DBS, ALS, KTT

Project administration: FRA, JAG Software: CRS, DBS, ALS

Supervision: FRA, JAG

Visualization: JMW, CRS, DBS

Writing – original draft: JMW, CRS, DBS

Writing – review & editing: ALS, KTT, FRA, JAG

## ACKNOWLEDGEMENTS

We thank all members of the Gagnon lab for discussions and comments. We thank Brian Dalley, Hailey Hollins, James Keener, Aaron McKenna, Alex Schier, and Kate Scuderi for feedback on the manuscript and technical assistance. We thank CZAR and CBRZ staff for zebrafish care, as well as the Center for High Performance Computing and the High-Throughput Genomics core facility, particularly Brett Milash and Martin Cuma. This work was supported by National Institutes of Health grants R35GM142950 (JAG), F30HD115391 (JMW), T32GM141848 (JMW and CRS),

T32HD007491 (ALS), and by National Science Foundation Graduate Research Fellowships (CRS and KTT).

## SUPPLEMENTAL FIGURES

**Supplemental Figure S1.**
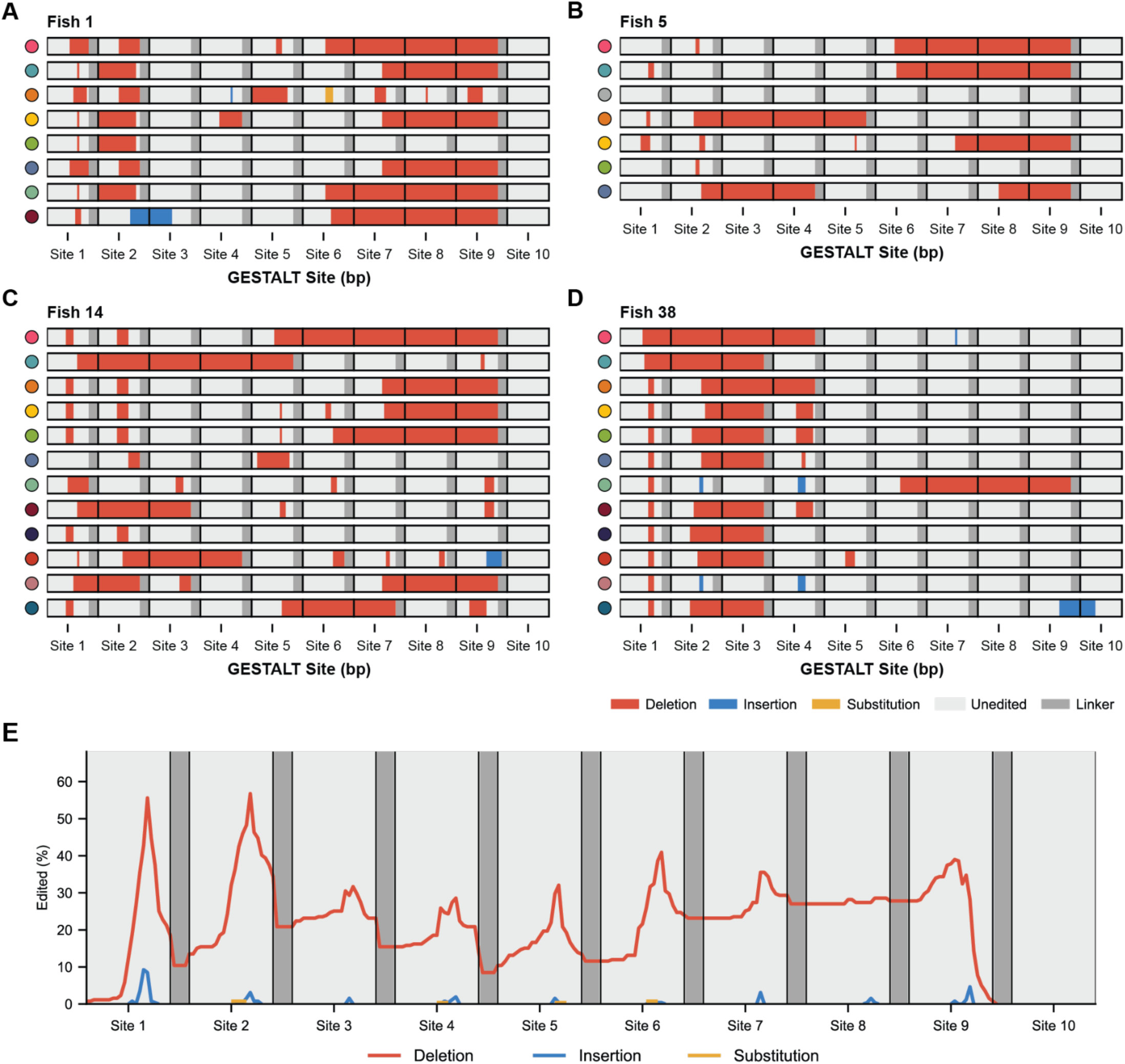
Maps of barcode edit locations. A-D) Diagrams that indicate the location of CRISPR edits on recovered barcodes from the first sperm sample of Fish 1, 5, 14, and 38. Dots on the left indicate the corresponding clone color in Fig. 1F. A legend specifying edit type is shown below panel D. E) A summary plot showing the cumulative locations of edits at each nucleotide position within CRISPR target sites averaged across all barcodes recovered from all samples from all fish.

**Supplemental Figure S2.**
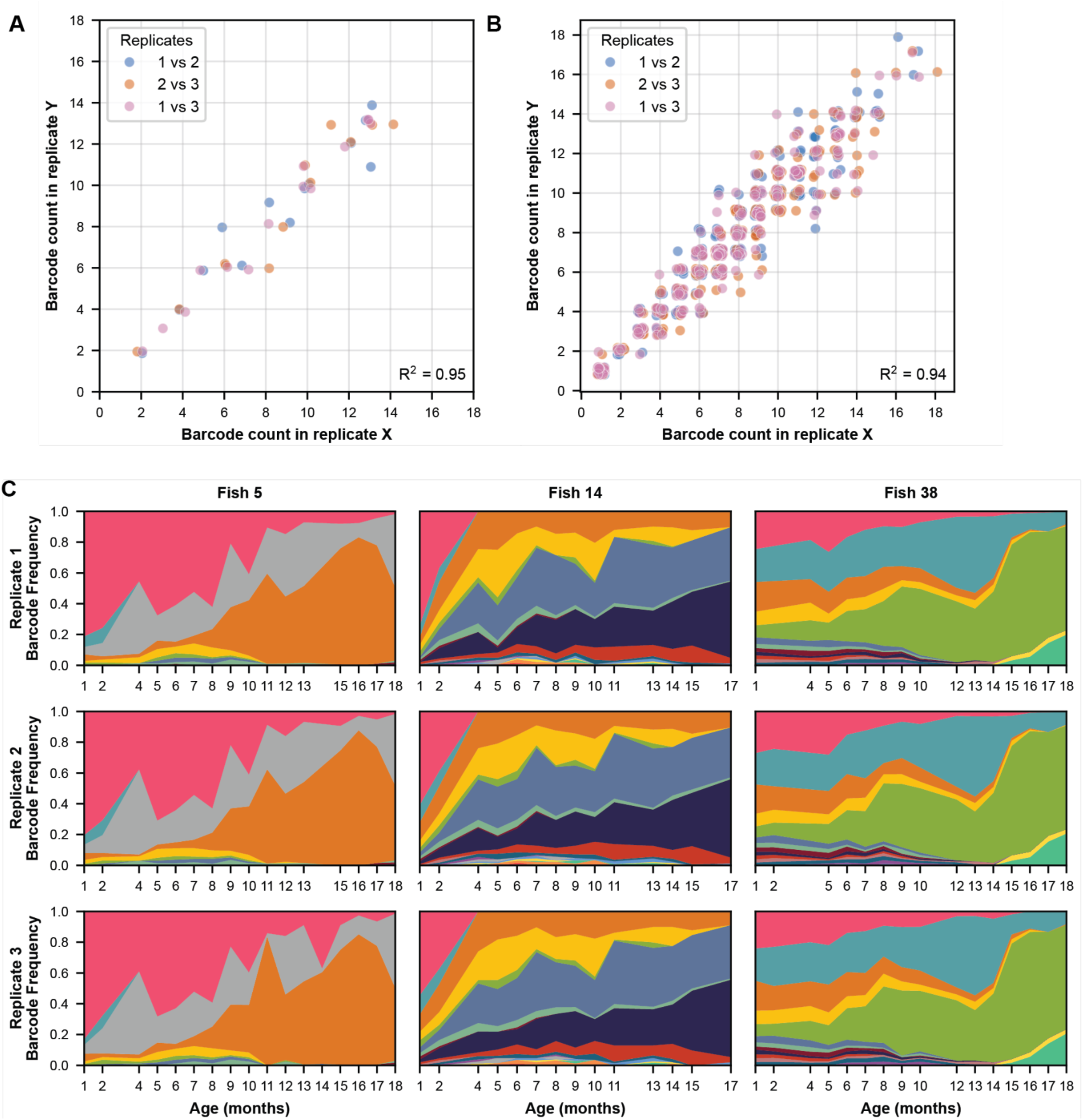
Technical replicates are consistent within samples. A) Pairwise comparison of barcode counts between replicates for all first-time point samples. Counts are always whole numbers but dots are jittered slightly to allow visualization. B) Pairwise comparison of barcode counts for all samples. C) Comparison of barcode frequencies across time between replicates for example fish.

**Supplemental Figure S3.**
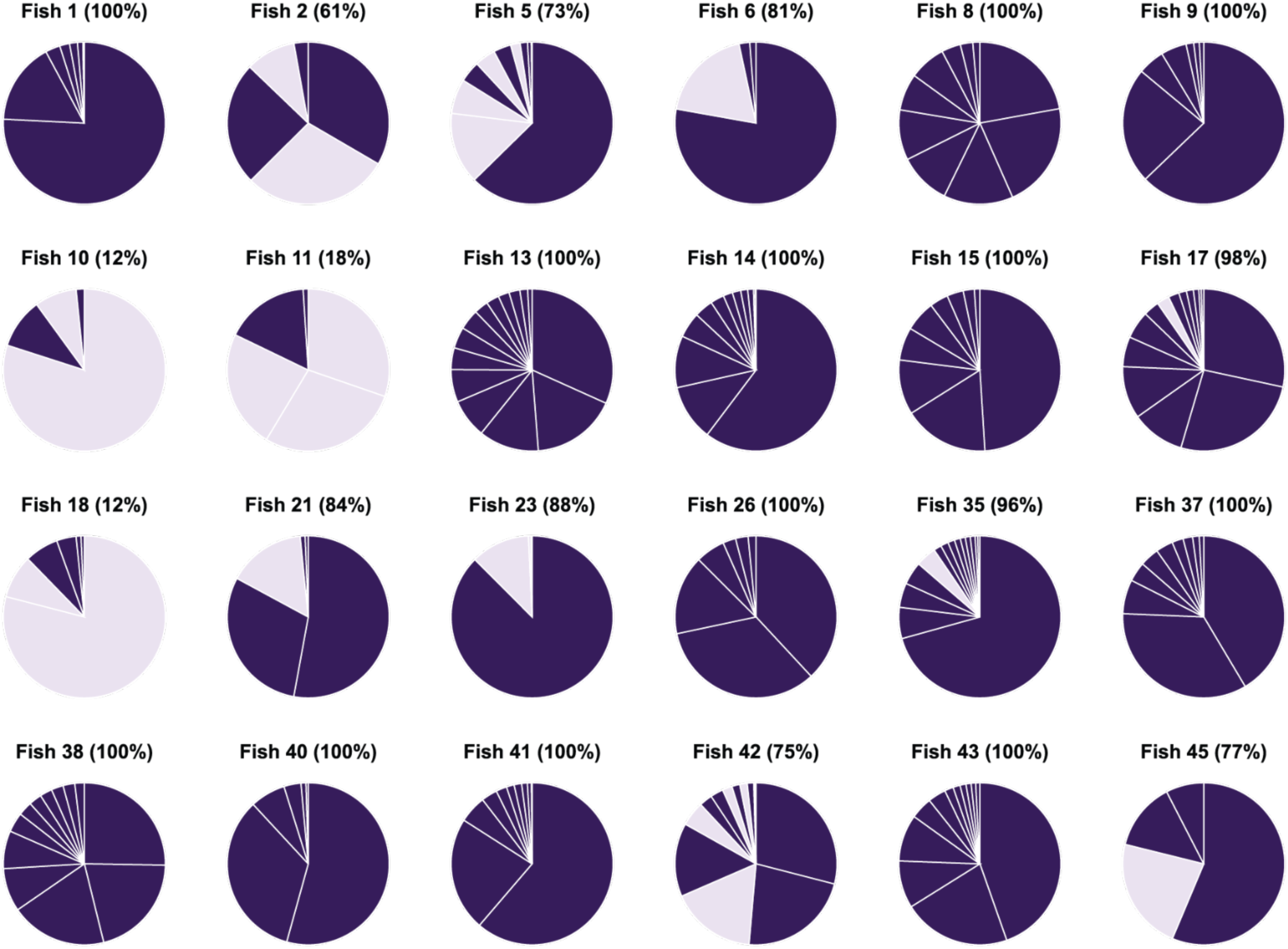
Unique barcodes recovered from each fish. Each slice represents a barcode collected from a fish in the first sample. Dark slices correspond to barcodes unique to that fish, while lighter barcodes were found in multiple fish. The percentage of reads that were from unique barcodes is given above each pie chart.

**Supplemental Figure S4.**
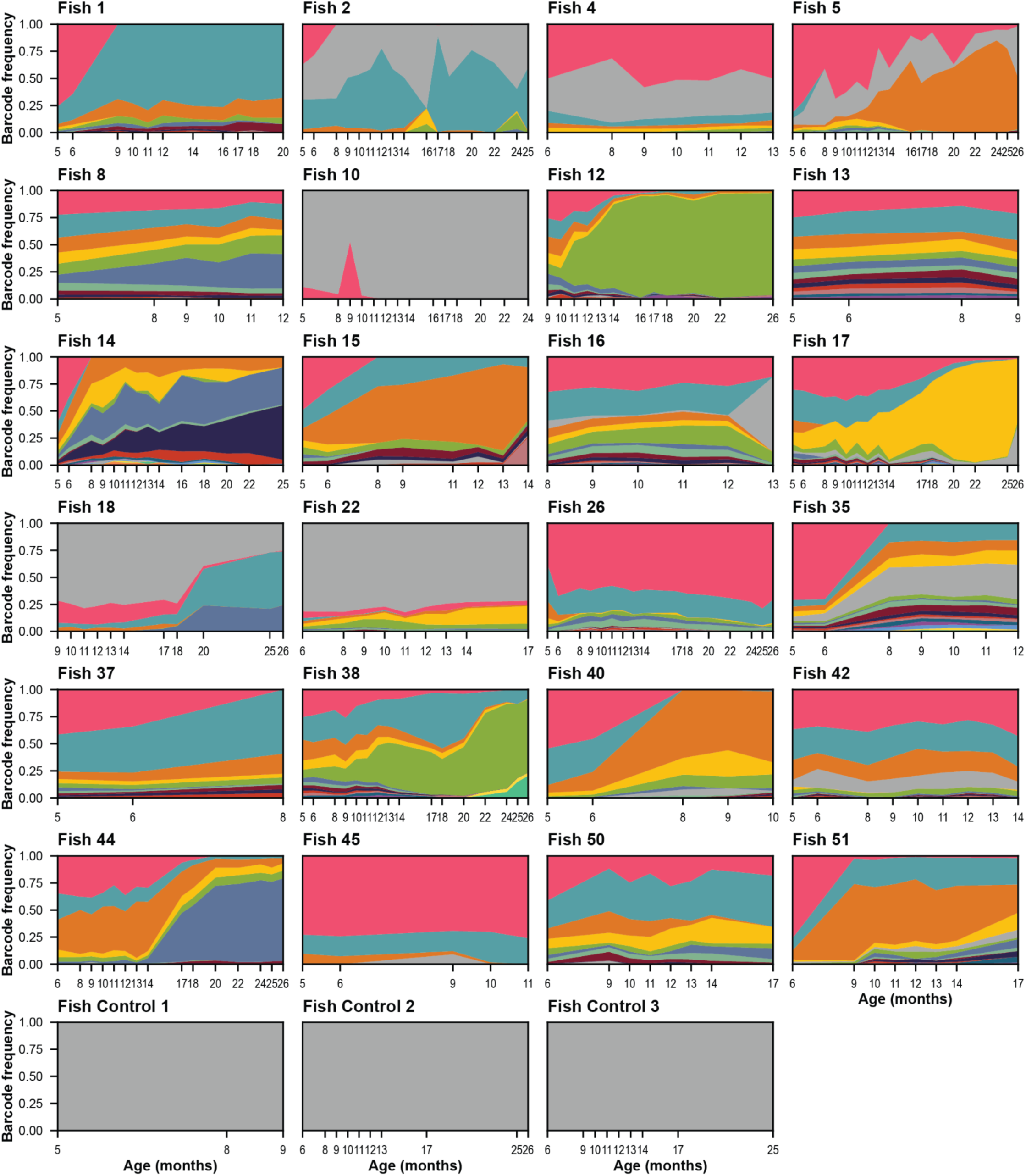
Barcode frequencies across time. Plotted here are all fish with sufficient data (see Methods). The same color scheme is used for each fish even though their barcodes are unique. Gray represents the unedited barcode. Control fish carry the barcode but did not undergo CRISPR-Cas9 editing.

**Supplemental Figure S5.**
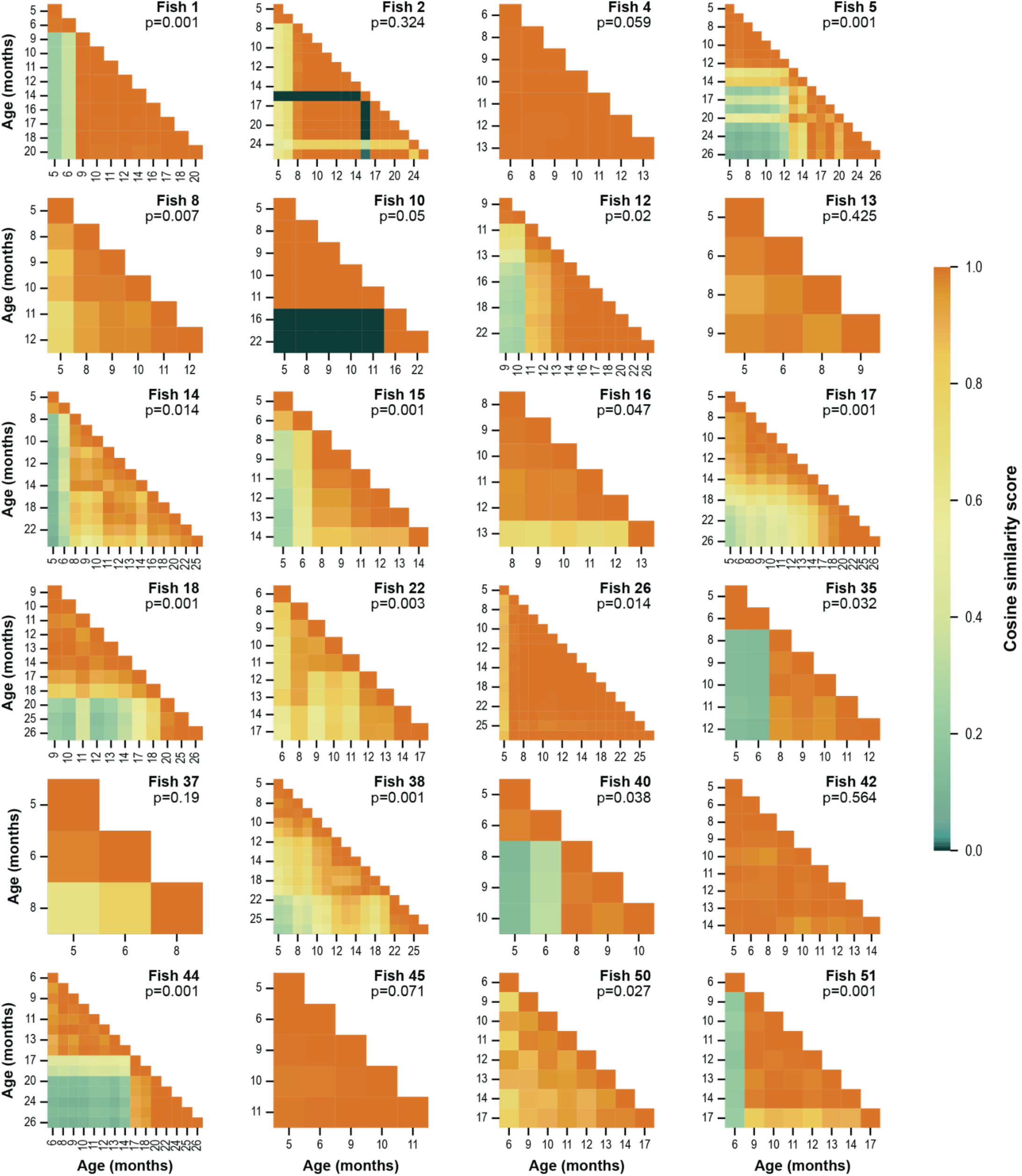
Heat maps comparing barcode distributions between samples for each fish. Color determined by cosine similarity score for each pairwise comparison. Diagonal in each facet represents comparison of that sample to itself (cosine similarity = 1). The Mantel test is applied to each fish to evaluate whether nearby samples are more similar than distant samples, with the p value shown in the top right of each plot (see Methods).

**Supplemental Figure S6.**
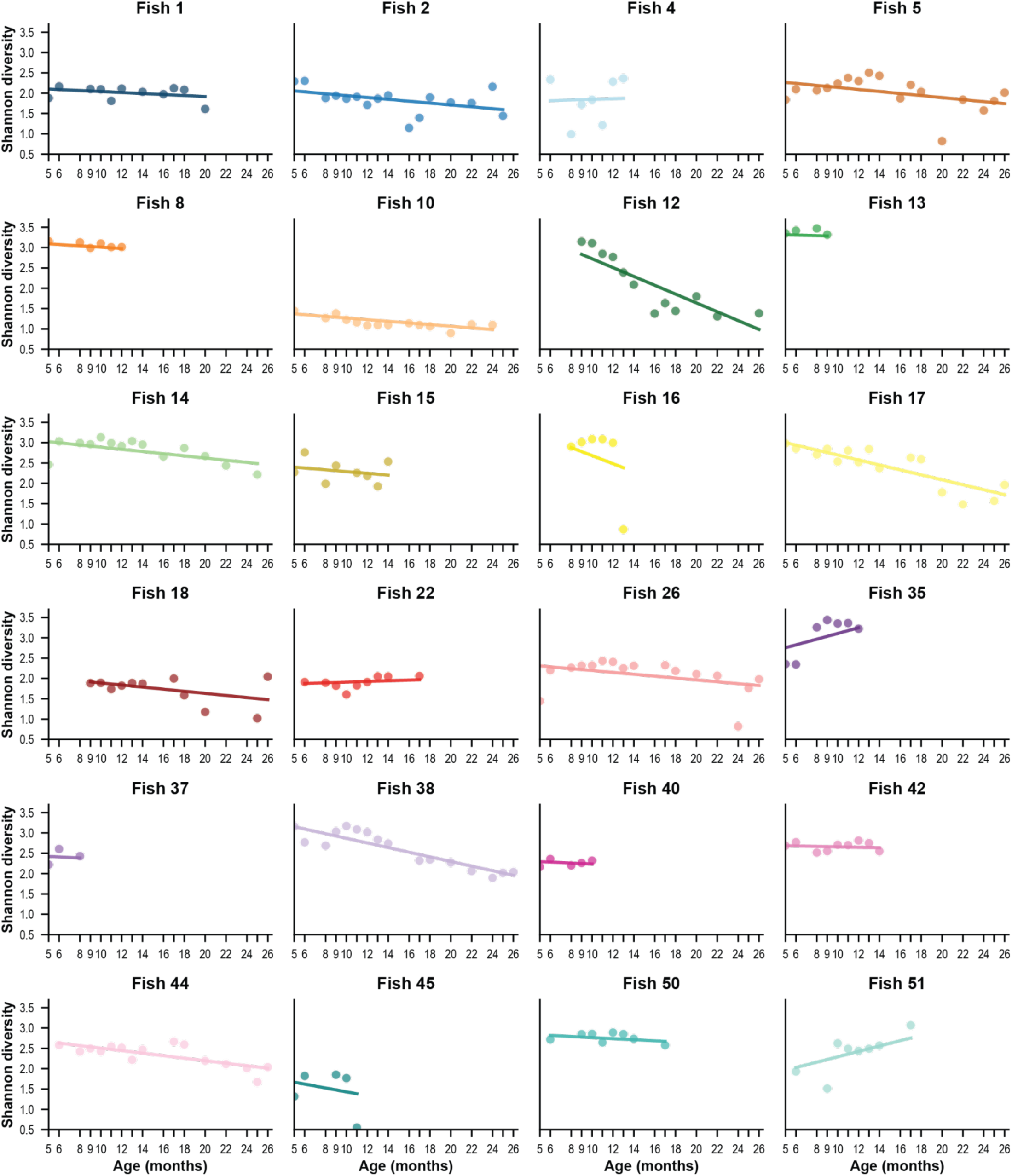
Shannon diversity index across time for each fish. Trendlines were fit using a linear mixed model.

**Supplemental Figure S7.**
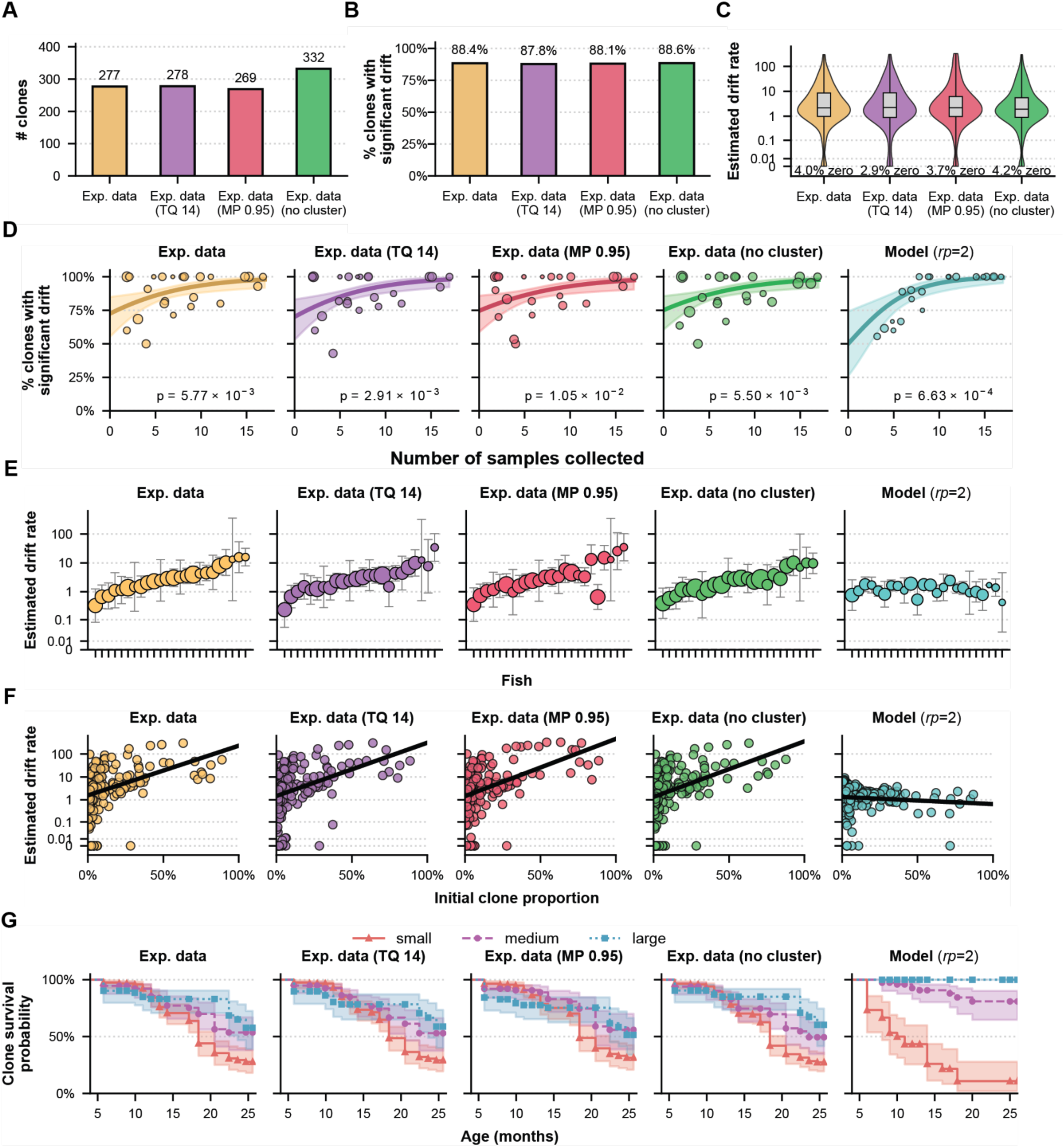
Features of clonal dynamics do not depend on data cleaning choices. The original data processing (yellow) is compared with three alternate methods. The first raises the GESTALT trim quality (TQ) parameter from 10 to 14 (purple), while the second instead sets the GESTALT match proportion (MP) parameter from 0.8 to 0.95 (red). The third alternative method skips the step that clusters barcodes based on Levenshtein distance (green). A) Total number of clones counted in all fish after different data cleaning pipelines. B) Extended version of Fig. 3D, showing the frequency of clones with significant evidence of drift. C) Extended version of Fig. 4A. D) Extended version of Fig. 3E. E) Extended version of Fig. 4B. F) Extended version of Fig. 4C. G) Extended version of Fig. 4D.

**Supplemental Figure S8.**
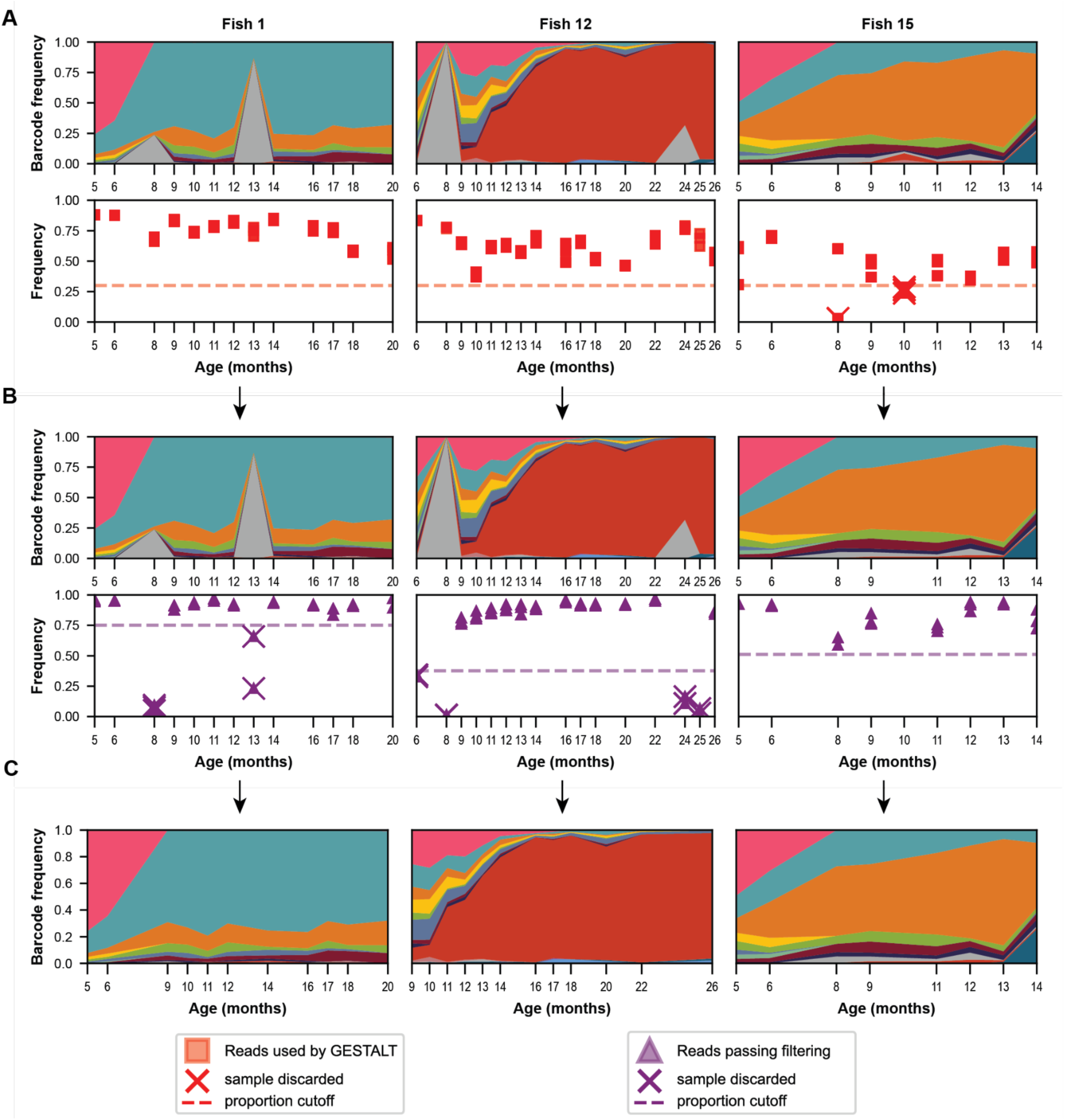
Anomaly detection in the quality control pipeline. Longitudinal data for fish 1, fish 12 and fish 15 are shown at each step of the anomaly filtering. A) The original data before anomaly detection. Proportion of reads used by GESTALT per sample are shown below. B) The data after removing samples with a low proportion of reads used by GESTALT. Proportion of reads remaining after the filtering pipeline are shown below. C) Samples with a low proportion of reads kept after the filtering pipeline are removed to produce the final longitudinal data. Note that the color of a barcode may change between panels depending on the number of removed barcodes.

**Supplemental Figure S9.**
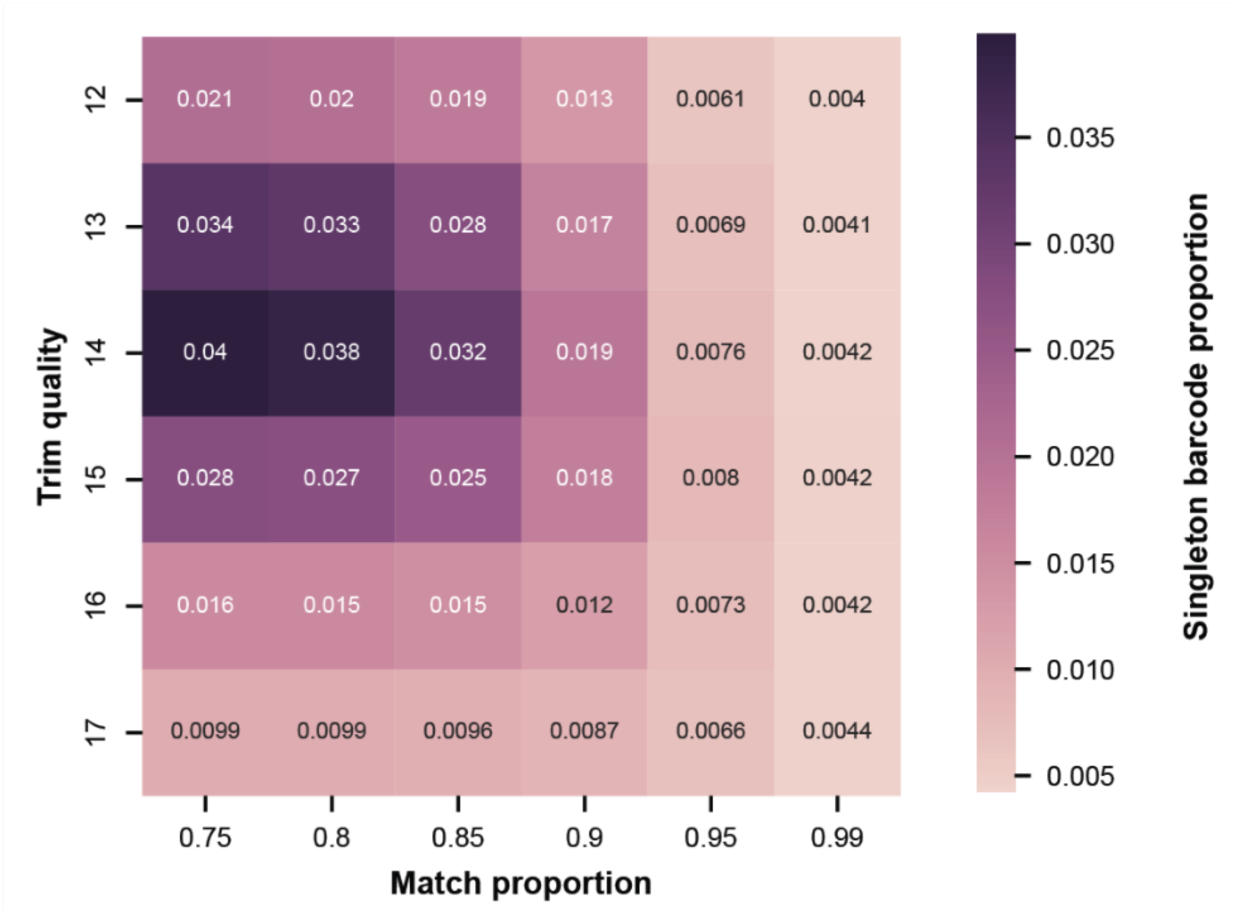
Contribution of GESTALT parameter choice in data processing. GESTALT takes several parameters that determine how raw reads are processed. Shown here is the frequency of singleton barcodes, which correspond to only one read in a fastq file, per combination of match proportion and trim quality parameters. These singleton barcodes are a result of error in sampling but are later filtered out by quality control. For this figure, GESTALT was run on a subset of all data (replicate 1 of sample 6 of seven fish).

**Supplemental Figure S10.**
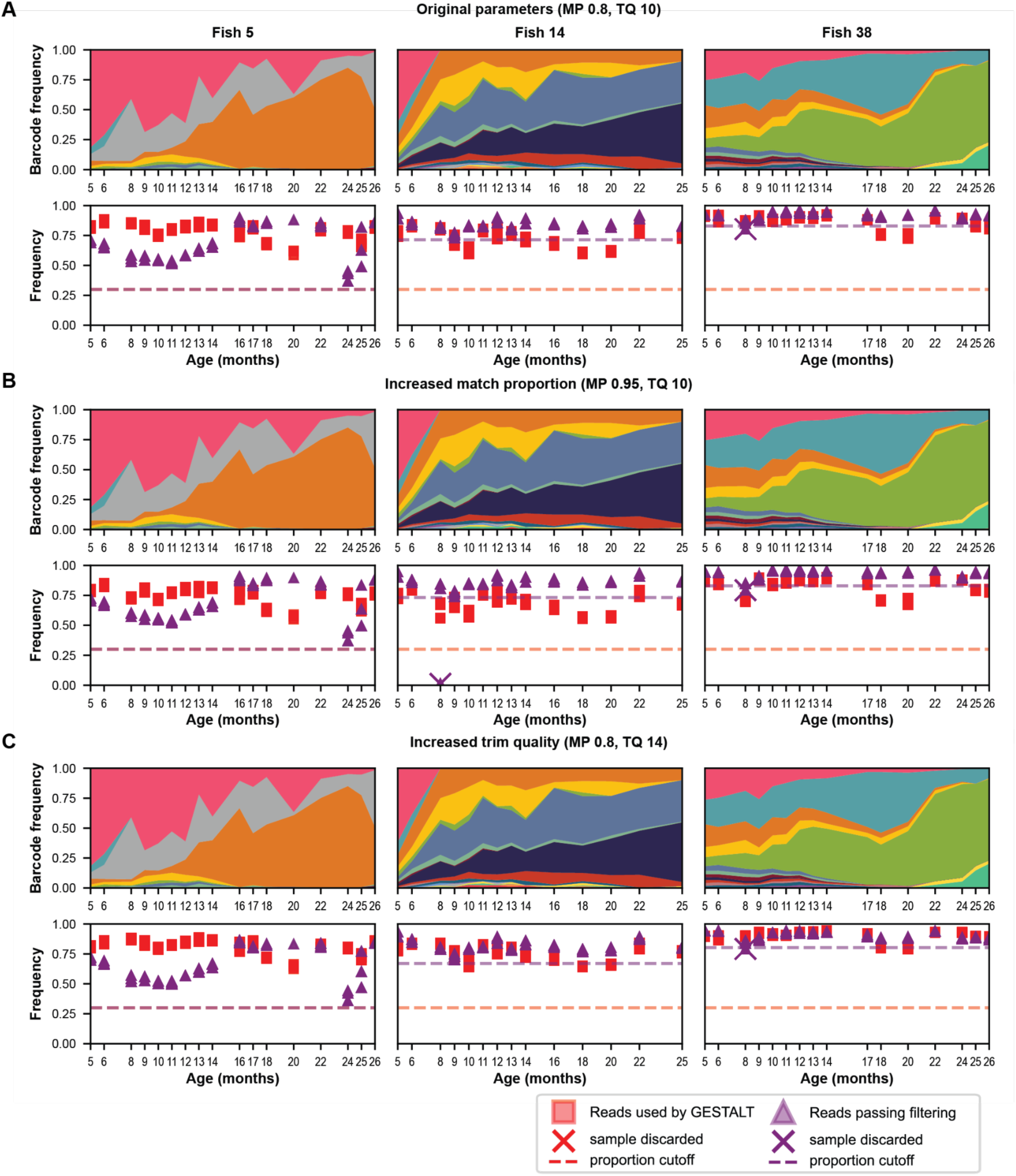
GESTALT output is not sensitive to pipeline parameter choices. To test the effect of GESTALT parameters on our findings, we re-ran our analysis using increased values for the trim quality and match proportion parameters. Longitudinal data is displayed for each set of parameters tested. A) The data obtained with the original parameters (match proportion = 0.8, trim quality = 10). B) The data obtained with match proportion increased to 0.95. C) The data obtained with trim quality increased to 14.

**Supplemental Figure S11.**
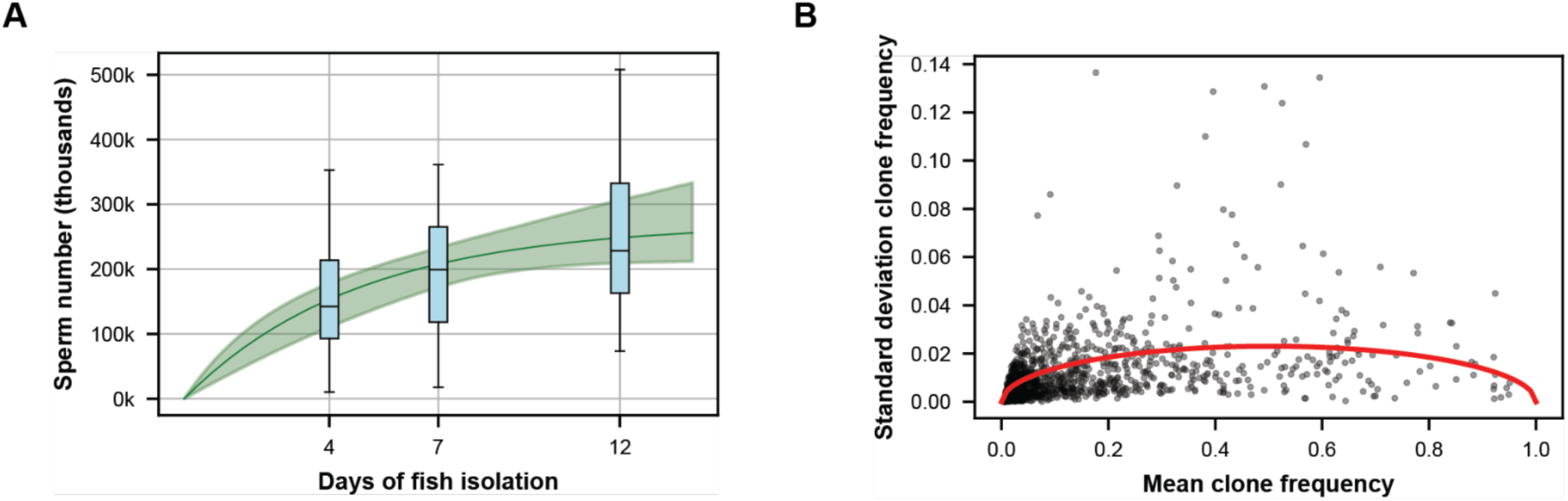
Estimation of model parameters. A) Selecting the sperm decay rate. The box plots show observed zebrafish sperm counts after different periods of isolation (Cattelan & Gasparini, 2021). We fit these data to a simple decay process, with the fit shown in green. The best fit gives a decay rate of δ=6.28 month^-1^. B) Estimate of the sperm pseudocount parameter. Each point corresponds to a clone in a single sample with more than one observed replicate after filtering. The *x* coordinate is the mean observed frequency of the clone and the y coordinate is the standard deviation across replicates. The red curve shows the expected standard deviation of a binomial sampling when choosing *M* = 470, which best fits the data using maximum likelihood.

## SUPPLEMENTARY INFORMATION

### Supplemental methods: model

We use a Moran process to describe spermatogonial stem cell differentiation and replacement in the testis (Moran, 1958). This continuous-time model was originally used to study the dynamics of allele frequencies in ecological populations, but here we use it to describe clonal frequencies in stem cell populations.

Assume that the total stem cell count is a constant *N*, with each stem cell assigned to an initial clone identity. Stem cells undergo spontaneous division at rate *r* month^-1^. Dividing cells can either symmetrically differentiate with probability *p*, or asymmetrically divide with probability 1–*p*. The neutral hypothesis assumes that all clones follow the same division rules, so *r* and *p* do not depend on clone identity. To maintain a constant population, we assume that when a cell differentiates, a random other cell (possibly from another clone) is selected to divide symmetrically. This process results in neutral drift because the size of one SSC clone has decreased by random chance, while another clone has increased in size. We therefore refer to this rate, *rp*, as the “strength of drift”. Asymmetric cell divisions do not create drift, but allow stem cells to produce sperm even when there is no drift.

When a stem cell differentiates, we assume that it produces 2*s_d_* sperm cells that inherit the clonal identity of the stem cell. Asymmetric divisions produce 2*s_d_* sperm cells. We assume that there is a time delay of *τ* months between stem cell differentiation and release of sperm into the lumen.

Sperm in the lumen decays according to a simple death process at rate δ month^-1^.

Sperm samples are simulated by removing all sperm residing in the lumen. Define *f_i_* to be the frequency of sperm in the total sperm population that are of clone *i*. Then recorded sperm data are taken as a multinomial sampling

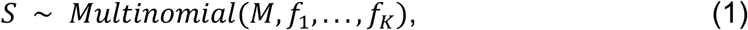

where *K* is the total number of clones. This choice assumes that the observed sperm are a random sample of the total sperm produced, with replacement. Because our experimental samples sequence a small subset of the total sperm collected, sampling with replacement is a suitable approximation.

The parameter *M* affects the amount of noise in each replicate, with larger values of *M* (more samples) being less noisy. We call the parameter the “pseudocount” of sperm. We choose its value so that the expected variance in observed sperm between simulated replicates matches the observed variance in the experimental replicates. This allows us to incorporate the other sources of noise present in our data, such as PCR bias. As a result, this parameter *M* will be less than the true number of sperm collected in experimental replicates.

### Parameter estimates

We choose *N*=1000 based on (Mullins et al., 1994) (estimates 500-1000 stem cells) and [Nobrega et. al] (counts at least 500 SSCs per fish). The parameter *r* is set to 36 cell cycles per month, which comes from (Nóbrega et al., 2010) estimating a cell cycle duration of 10–30 (mean 20) hours. Because differentiations induce symmetric divisions in our model, the effective number of cell cycles in our simulations is larger than 36. However, we found that for a fixed value of *rp*, the value of *r* does not significantly affect results, so we kept *r*=36.

The time delay *τ* from SSC to spermatid is 6 days, or 0.2 months (Leal et al., 2009). The number of barcoded sperm is set to *s_d_*= 631, which is half the number of spermatids found in a single Sertoli cyst (Leal et al., 2009) (the other half of sperm do not have a barcode since they are haploid).

The sperm pseudocount *M* is chosen using maximum likelihood to best match the variance observed in the experimental replicates. We find that *M*=470 best matches the noise found across replicates (Fig. S11B). The sperm decay rate δ is fit using data from (Cattelan & Gasparini, 2021), who count the sperm cells sampled from a fish 4, 7, and 12 days after the previous sample. We fit their data to a simple model that assumes sperm are produced at constant rate *C* (sperm per month) and decays at rate δ (deaths per month). By fitting their data to this model using scipy’s *curve_fit*, we find that the best fit parameters are *C*=1,698,090 and δ=6.28. The 95% confidence interval for δ is 3.93-8.63. Table (S1) summarizes the model parameters.

**Supplementary Table S1.**
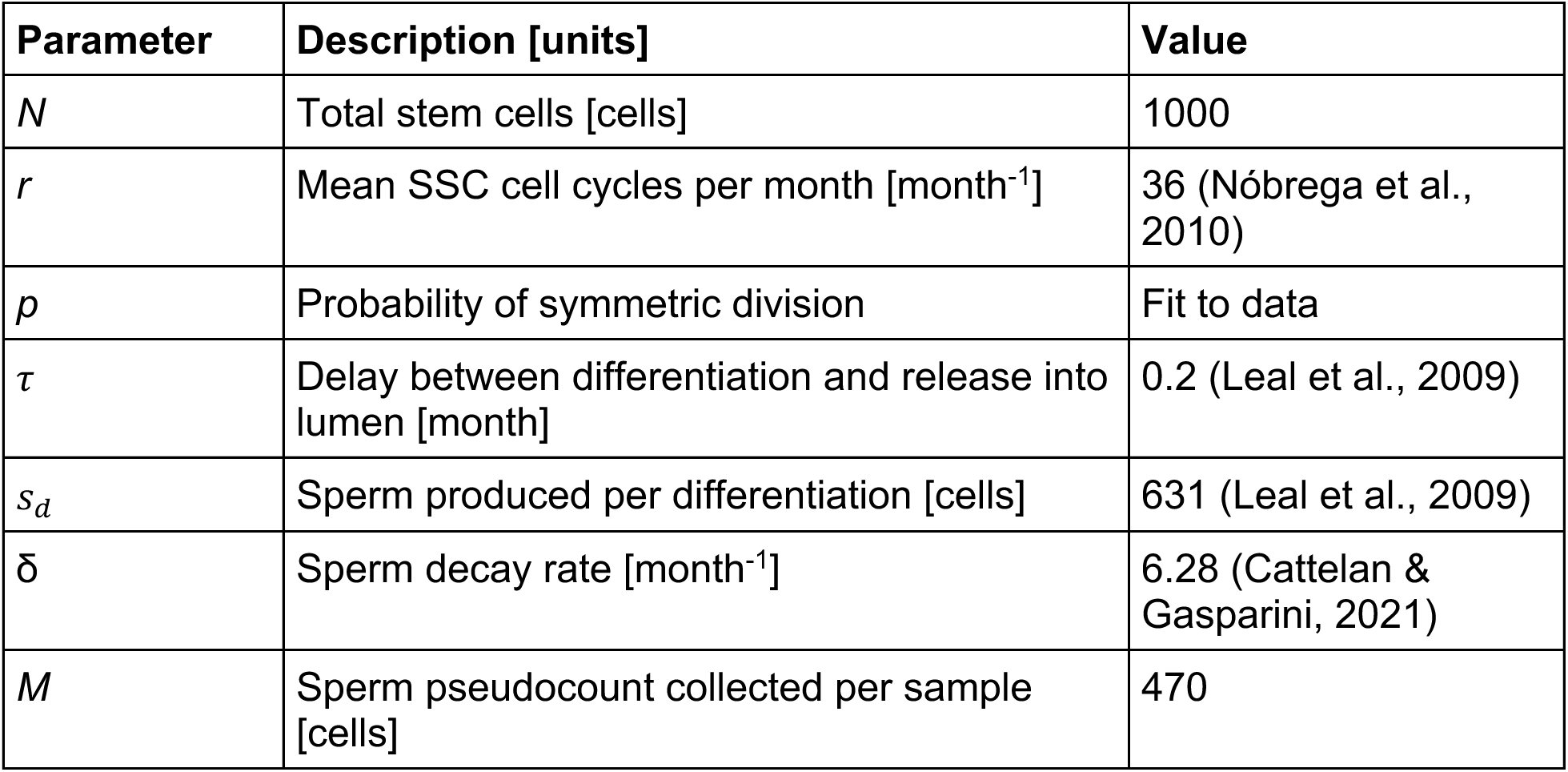
Mathematical model parameters.

| Parameter | Description [units] | Value |
| --- | --- | --- |
| $N$ | Total stem cells [cells] | 1000 |
| $r$ | Mean SSC cell cycles per month [month<sup>-1</sup>] | 36 (Nóbrega et al., 2010) |
| $p$ | Probability of symmetric division | Fit to data |
| $\tau$ | Delay between differentiation and release into lumen [month] | 0.2 (Leal et al., 2009) |
| $s_d$ | Sperm produced per differentiation [cells] | 631 (Leal et al., 2009) |
| $\delta$ | Sperm decay rate [month<sup>-1</sup>] | 6.28 (Cattelan & Gasparini, 2021) |
| $M$ | Sperm pseudocount collected per sample [cells] | 470 |

### Estimating drift rate

We define the “strength of drift” of a clone to be the value of *rp* in the model that is most likely to reproduce that clone’s sampling data. This allows us to quantify the amount of drift seen in our experimental data using the model. Because the model assumes clones are neutral, meaning that all clones follow the same rules, we can measure the rate of drift in a single clone using only its frequency data relative to the sum of all remaining clones. This reduces the system to the two-clone case.

The PCR “reads” of each clone are not comparable between samples because PCR efficiency varies between samples. Instead, we convert the observed clone proportions to sperm pseudocounts defined earlier. Let *O_i_* denote the observed sperm pseudocounts of a target clone at sampling time *t_i_*. Then the likelihood function is defined as the probability of observing all of the collected sperm data given a choice of *rp*:

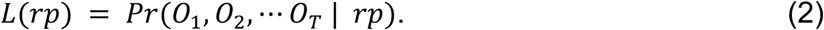

In principle, computing *L*(*rp*) would require simulating the time evolution of the joint probability distribution of three variables: the number of stem cells in the target clone, the sperm produced by that clone, and the sperm produced by remaining clones. Because the cell count for each of these variables is on the order of hundreds or more, this is computationally infeasible.

To overcome this challenge, we introduce a few simplifying assumptions:

- Rather than tracking the joint distribution of stem cell population and sperm produced by the target clone, we instead track only the distribution of stem cell populations under the classical Moran process.
- The sperm produced by a clone is assumed to be an instantaneous sample of the stem cell population *τ* months prior, rather than tracking the accumulation of sperm over an interval of time. In other words, rather than sampling from accumulated sperm frequencies as *f_i_* in Equation (1), we set *f_i_* to be the stem cell frequencies.

With these assumptions, our model simplifies to a first-order Hidden Markov model, which is much more suitable for inference (Rabiner & Juang, 1986).

Between two sampling times *t_i_* and *t_i_*_+1_, we specify a transition probability matrix *T*, where *T*[*i*, *j*] is the probability that the stem cell population changes from *i* to *j* in time (*t_i_*_+1_ − *t_i_*). This matrix is readily computed from the Moran process at rate *rp* (Karlin, 2014). We also define an emission matrix *E*, where *E*[*i*, *j*] is the probability of observing *j* sperm given the stem cell population equals *i* at the current sample given by equation (1). The likelihood function is then evaluated using the *forward algorithm* (also known as *expectation maximization*), which uses dynamic programming to compute the likelihood (Equation 2) in *O*(*TN*^2^*M*) time, where *T* is the number of samples of a given clone (Rabiner & Juang, 1986).

We define the strength of drift of a clone to be the choice of *rp* that maximizes this likelihood,

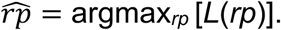

This method also allows us to construct confidence intervals for the drift rate *rp*. We set the lower and upper limits of this interval to be values of *rp* such that

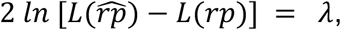

where λ is a chosen likelihood ratio statistic. We choose λ=3.84, which corresponds to a 95% confidence interval (Zar, 2014).

Our method differs from other algorithms that estimate the *effective population size N_e_* as a measure of drift strength. (Williamson & Slatkin, 1999), for instance, use a hidden Markov Model and maximum likelihood to estimate effective population size *N_e_* from sampled allele frequency. Our method estimates a continuous-valued drift rate *rp* rather than an integer-valued population size *N_e_*. This improves runtime since our assumed cell population is held constant, and allows us to use gradient-based optimization methods. Other approaches use variance-based methods to estimate effective population size (Tataru et al., 2017). However, we choose this likelihood-based approach because it connects more naturally to our original (unsimplified) model and allows us to more directly design hypothesis tests and confidence intervals.

### Estimator bias

Our maximum likelihood estimator could be biased, meaning that it tends to over- or under-estimate the drift strength *rp*. This is especially relevant because the model we use for parameter estimation is not the same as the full model that we simulate. To evaluate the robustness of our inference method, we estimate *rp* in simulated datasets from the full model. We simulated 240 fish per value of *rp* (10 simulated fish per experimental fish). Supplementary Table S2 summarizes the bias of our method. We find that for most true values of *rp*, our method underestimates values by 5-10%. The confidence intervals around each estimate contain the correct value of *rp* approximately 94% of the time across all test cases. This indicates that the model simplifications do not limit the effectiveness of our inference.

**Supplementary Table S2.** Bias and variance of maximum likelihood estimator. For each underlying drift rate *rp*, 240 clones were simulated. The mean estimate for *rp* and its relative bias are shown. The percentage of confidence intervals (CIs) that contain the ground truth drift rate are also shown.

| True $rp$ | Mean ± SD estimate | Relative bias (%) | % CIs containing true $rp$ |
| --- | --- | --- | --- |
| 0.0 | 0.06 ± 0.19 | NA | 97.73% |
| 0.5 | 0.53 ± 0.55 | 5.47% | 95.37% |
| 2.0 | 1.88 ± 1.45 | -5.89% | 94.64% |
| 5.0 | 4.75 ± 3.24 | -5.09% | 93.82% |
| 8.0 | 7.38 ± 4.70 | -7.73% | 93.36% |
| 15.0 | 13.44 ± 9.00 | -10.38% | 93.30% |
| 25.0 | 22.92 ± 14.81 | -8.32% | 93.36% |

